# Presynaptic mitochondria volume and abundance increase during development of a high-fidelity synapse

**DOI:** 10.1101/689653

**Authors:** Connon I. Thomas, Christian Keine, Satoko Okayama, Rachel Satterfield, Morgan Musgrove, Debbie Guerrero-Given, Naomi Kamasawa, Samuel M. Young

## Abstract

The calyx of Held, a large glutamatergic presynaptic terminal in the auditory brainstem undergoes developmental changes to support the high action-potential firing rates required for auditory information encoding. In addition, calyx terminals are morphologically diverse which impacts vesicle release properties and synaptic plasticity. Mitochondria influence synaptic plasticity through calcium buffering and are crucial for providing the energy required for synaptic transmission. Therefore, it has been postulated that mitochondrial levels increase during development and contribute to the morphological-functional diversity in the mature calyx. However, the developmental profile of mitochondrial volumes and subsynaptic distribution at the calyx of Held remains unclear. To provide insight on this, we developed a helper-dependent adenoviral vector (HdAd) that expresses the genetically encoded peroxidase marker for mitochondria, mito-APEX2, at the mouse calyx of Held. We developed protocols to detect labeled mitochondria for use with serial block face scanning electron microscopy to carry out semi-automated segmentation of mitochondria, high-throughput whole terminal reconstruction and presynaptic ultrastructure in mice of either sex. Subsequently, we measured mitochondrial volumes and subsynaptic distributions at the immature postnatal day 7 (P7) and the mature (P21) calyx. We found an increase of mitochondria volumes in terminals and axons from P7 to P21 but did not observe differences between stalk and swelling subcompartments in the mature calyx. Based on these findings, we propose that mitochondrial volumes and synaptic localization developmentally increase to support high firing rates required in the initial stages of auditory information processing.

**Significance Statement:** Elucidating the developmental processes of auditory brainstem presynaptic terminals is critical to understanding auditory information encoding. Additionally, morphological-functional diversity at these terminals is proposed to enhance coding capacity. Mitochondria provide energy for synaptic transmission and can buffer calcium, impacting synaptic plasticity; however, their developmental profile to ultimately support the energetic demands of synapses following the onset of hearing remains unknown. Therefore, we created a helper-dependent adenoviral vector with the mitochondria-targeting peroxidase mito-APEX2 and expressed it at the mouse calyx of Held. Volumetric reconstructions of serial block face electron microscopy data of immature and mature labeled calyces reveal that mitochondrial volumes are increased to support high firing rates upon maturity.

## Introduction

During development of auditory brainstem circuits, many synapses undergo morphological and physiological transformations to support the high action-potential (AP) firing rates required in the initial stages of auditory information processing (Yu and Goodrich, 2014). Synaptic transmission and developmental remodeling are energetically demanding processes (Harris et al., 2012), and mitochondria are essential organelles for supplying energy for these processes (Devine and Kittler, 2018). In addition, mitochondria play a role in calcium (Ca^2+^) buffering which can impact synaptic plasticity (Vos et al., 2010). Therefore, understanding how presynaptic mitochondria volume and distribution change with development is integral to understanding how high-fidelity AP firing is achieved and maintained in the auditory brainstem.

The calyx of Held, a large glutamatergic presynaptic terminal in the medial nucleus of the trapezoid body (MNTB) in the auditory brainstem undergoes considerable morphological and physiological transformations around the onset of hearing at the end of the second postnatal week (Baydyuk et al., 2016). At the physiological level, the calyx of Held/MNTB synapse shifts from exhibiting frequent synaptic failures before hearing onset to reliable synaptic transmission at firing rates up to several hundred Hertz (Taschenberger and von Gersdorff, 2000). In parallel there is a reduction in release probability (*Pr*) (Taschenberger and von Gersdorff, 2000; Iwasaki and Takahashi, 2001) and a shift from micro-to nano-domain release due to tighter coupling—the distance between synaptic vesicles (SVs) and voltage gated calcium channels (VGCCs) (Fedchyshyn and Wang, 2005; Wang et al., 2008; Chen et al., 2015). At the morphological level, the calyx transforms from cup-like to a highly fenestrated structure consisting of stalks and swellings (Kandler and Friauf, 1993; Rowland et al., 2000; Wimmer et al., 2006), and undergoes ultrastructural changes such as smaller active zones (AZs) and a reduced number of docked SVs (Taschenberger et al., 2002). Finally, the mature calyx contains mitochondria tethered to the presynaptic plasma membrane (Rowland et al., 2000; Perkins et al., 2010) and SVs are tightly packed around mitochondria (Wimmer et al., 2006).

In addition to these developmental changes, a morphological-functional diversity at the mature calyx has been shown, in which calyces with many swellings have lower synaptic depression and SV release probability (Grande and Wang, 2011). Furthermore, stalks and swellings differ in Ca^2+^ buffering and SV coupling states, with stalks having tighter SV coupling than swellings (Fekete et al., 2019). Nanodomain release corresponds to tight coupling with SV-VGCC distances of ∼25 nm and is considered to be energetically favorable (Eggermann et al., 2012); therefore, the energetic demands of AZs at stalks are predicted to be lower than swellings. However, the developmental changes in mitochondria volumes and subsynaptic distribution between stalks and swellings in the calyx remain unknown. To address these questions, we carried out volumetric reconstructions at the electron microscopic (EM) level of calyces and mitochondria at the immature postnatal day 7 (P7) and mature P21 calyx. To do so, we created a helper-dependent adenoviral vector (HdAd) that expresses the mitochondrial EM marker mito-APEX2 (Martell et al., 2012; Lam et al., 2015; Martell et al., 2017). Since serial block face scanning EM (SBF-SEM) makes large scale reconstructions more accessible than transmission EM (TEM) (Lichtman and Denk, 2011), we developed protocols permitting identification of mito-APEX2 with SBF-SEM and reconstructed multiple mito-APEX2 positive terminals while resolving presynaptic ultrastructure. We found a developmental increase in mitochondrial volumes in the calyx and its respective axon while mitochondrial volumes were not different between stalk and swellings of the adult calyx. These data suggest, that mitochondrial volumes developmentally increase to support high firing rates and may contribute to morphological-functional diversity at the calyx. Finally, since mito-APEX2 permits identification of calyces while preserving ultrastructure, we optimized protocols using automated tape-collecting ultramicrotome serial section SEM (ATUM ssSEM) to analyze single-AZ ultrastructure in partially reconstructed calyces. We propose our viral vectors and methods will have broad applications for quantitative analyses of subsynaptic organization of SVs, mitochondria, and their morphological continuum using TEM, ssSEM and SBF-SEM.

## Materials and Methods

### Animals

All experiments were performed in accordance with animal welfare laws and approved by the Institutional Committee for Care and Use of Animals at the Max Planck Florida Institute for Neuroscience and the University of Iowa and complied with accepted ethical best practice. Animals were housed at a 12-hour light/dark cycle and had access to food and water ad libitum. Experiments were performed on C57Bl/6J mice (RRID:IMSR_JAX:000664, The Jackson Laboratory, Bar Harbor, USA) of either sex. Virus constructs were injected at postnatal day 1 (P1) and experiments were performed at P7 (immature calyces) or P21 (functionally mature calyces).

### DNA construct and recombinant viral vector production

Mito-V5-APEX2 (Lam et al., 2015), a gift from Alice Ting (Addgene plasmid #72480, http://n2t.net/addgene:72480, RRID:Addgene_72480), was cloned into the EcoRI and NotI sites of the synapsin expression cassette. This cassette included the 470 bp human synapsin (hsyn) promoter, the minute virus of mice (mvm) intron, and the bovine growth hormone polyA sequence (Montesinos et al., 2011). HdAd was produced as previously described (Montesinos et al., 2016). The mito V5 expression cassette was cloned into the AscI site of a modified version of pdelta 28E4, gift from Dr. Philip Ng, using InFusion (Clontech). This version of pdelta 28E4 was modified to also contain a separate neurospecific EGFP expression cassette driven by the hsyn promoter (Lubbert et al., 2017). HdAd was produced as previously described (Montesinos et al., 2016). Briefly, pHdAd mito-APEX2 hsyn EGFP was digested with PmeI and into 116 producer cells. HdAd was serially amplified in five consecutive passages. Each successive passage was performed after cytopathic effect occurred and cell lysates were subjected to three freeze-thaw cycles to lyse cells and thereby release the viral particles. HdAd was stored at −80°C in storage buffer containing 10 mM HEPES, 250 mM sucrose, and 1 mM MgCl_2_ at pH 7.4.

### Virus injections

Virus injections at P1 were performed as described previously (Chen et al., 2013). Briefly, mice were anesthetized by hypothermia for 5 min in an ice bath and a stereotactic injection of ∼1 μL of HdAd (total number of virus particles <1e9) syn mito-V5-APEX2 syn EGFP into the cochlear nucleus was performed at a rate of 1 μL/min using glass pipettes (Blaubrand, IntraMARK). Following the injection, the glass needle was left in place for 1 min to dissipate the pressure and then slowly retracted. Animals were then placed under an infrared heat lamp and allowed to recover before being returned to their respective cages with their mother.

### Acute brain slice preparation

Acute coronal brainstem slices (200 μm) containing the medial nucleus of the trapezoid body (MNTB) were prepared from control and EGFP-mito-APEX2-injected animals using a Leica VT 1200S vibratome equipped with zirconia ceramic blades (EF-INZ10, Cadence Blades) in low-calcium artificial cerebrospinal fluid (aCSF) solution, containing (in mM): 125 NaCl, 2.5 KCl, 3 MgCl_2_, 0.1 CaCl_2_, 10 glucose, 25 NaHCO_3_, 1.25 NaH_2_PO_4_, 0.4 L-ascorbic acid, 3 myo-inositol, and 2 Na-pyruvate, pH 7.3–7.4. The blade was advanced at a speed of 20-50 μm/s. Slices were immediately transferred to an incubation beaker containing standard extracellular solution (same as above but using 1 mM MgCl_2_ and 1.2 mM CaCl_2_ at 37°C, continuously bubbled with 95% O_2_– 5% CO_2_). After approximately 45 min of incubation, slices were transferred to a recording chamber with the same saline at room temperature (∼22°C).

### Electrophysiology

Fiber stimulation was performed as described previously (Dong et al., 2018). Briefly, to record AP-evoked EPSCs, fibers of the trapezoid body were stimulated using a bipolar platinum-iridium electrode (FHC, Model MX214EP) positioned medially of the MNTB. Postsynaptic MNTB neurons were whole-cell voltage clamped at −73 mV using an EPC10/2 amplifier controlled by Patchmaster Software (version 2×90.2, HEKA Elektronik, RRID:SCR_000034). Slices were continuously perfused with standard aCSF solution and visualized by an upright microscope (BX51WI, Olympus) through a 60x water-immersion objective (LUMPlanFL N, Olympus) and an EMCCD camera (Andor Luca S, Oxford Instruments). To identify calyces transduced with the EGFP-mito-APEX2 virus, the slice was illuminated at an excitation wavelength of 480 nm using a Polychrome V xenon bulb monochromator (TILL Photonics). During recordings, the standard extracellular solution was supplemented with 1 mM kynurenic acid (Tocris, Cat# 0223) to avoid desensitization and saturation of postsynaptic AMPA receptors, 50 μM D-AP-5 (Tocris, Cat# 0106) to block NMDA receptors, and 20 μM bicuculline (Tocris, Cat# 0131) and 5 μM strychnine (Tocris, Cat# 2785) to block inhibitory GABA- and glycine receptors, respectively. Patch pipettes were fabricated using a PIP6 (HEKA) and had a resistance of ∼3–4 MΩ when filled with the following (in mM): 130 Cs-gluconate, 20 tetraethylammonium (TEA)-Cl, 10 HEPES, 5 Na_2_-phosphocreatine, 4 MgATP, 0.3 NaGTP, 6 QX-314, and 5 EGTA, pH 7.2 (315 mOsm). Liquid junction potential was 13 mV and corrected for all reported voltages. Data were acquired at a sampling rate of 50 kHz. Series resistance (3–8 MΩ) was compensated online to <3.0 MΩ, and the remaining series resistance was further compensated offline to 0 MΩ with a time lag of 20 μs (Traynelis, 1998) using a custom-written MATLAB function (version 9.4; Mathworks, RRID:SCR_001622).

### Electrophysiological data analysis

Electrophysiological data were imported to MATLAB using a custom-modified version of sigTool (Lidierth, 2009) and analyzed offline with custom-written MATLAB functions. EPSC amplitudes were measured as peak amplitude minus baseline preceding the EPSC. The RRP size was calculated using the back-extrapolation method as described earlier (Neher, 2015).

### EM slice preparation

Slices for EM were prepared independently of acute slices for electrophysiology experiments. P7 or P21 animals previously injected with the EGFP-mito-APEX2 viral vector were anesthetized with an intraperitoneal injection of tribromoethanol (250 mg/ kg of body weight) and perfused transcardially with 0.9% NaCl in MilliQ water, followed by a fixative containing 2% paraformaldehyde and 1-2% glutaraldehyde in 0.1 M cacodylate buffer (CB, pH 7.4) containing 2 mM CaCl_2_. Brains were extracted and post fixed in the same fixative for 2 hours, then washed in CB. The brainstem was sliced coronally at a thickness of 50 µm (TEM, ssSEM) or 100 µm (SBF-SEM) using a Leica VT 1200S vibratome in 0.1 M CB. The EGFP expression at the injection site and in the contralateral MNTB was confirmed under an epifluorescence microscope (CKX41, Olympus). All EM images of calyx terminals were taken from the middle region of the MNTB.

### TEM sample processing and data analysis

Slices for conventional TEM imaging were reacted in a solution containing 0.05% DAB (3,3’-Diaminobenzidine, Sigma Aldrich) in CB (pH 7.4) for 1 hour on ice. Slices were then reacted with 0.03% H_2_O_2_ (v/v) in the same well for 5-10 minutes and rinsed thoroughly. Labeling was confirmed using a standard brightfield microscope. Tissue was reacted with 1% aqueous osmium tetroxide for 1 hour on ice, rinsed with distilled water, reacted with 1% aqueous uranyl acetate for 30 minutes in the dark at 4°C, and rinsed again with distilled water. Tissue was then dehydrated in a graded ethanol series (30, 50, 70, 90, 100%), 1:1 ethanol to acetone, 100% acetone, 1:1 acetone to propylene oxide (PO), and 100% PO, each 2 × 5 minutes. Tissue was infiltrated using 3:1 PO to Durcupan resin (Sigma Aldrich) for 2 hours, 1:1 PO to resin for 2 hours, and 1:3 PO to resin overnight, then flat embedded in 100% resin on a glass slide and covered with an Aclar sheet at 60°C for 2 days. The MNTB region contralateral to the injected cochlear nucleus was trimmed, mounted on empty resin blocks with cyanoacrylate glue (Krazy Glue, Elmer’s Products), and sectioned at 40 nm using a Leica UC7 ultramicrotome. Sections were counterstained with uranyl acetate and lead citrate, and examined in a Tecnai G2 Spirit BioTwin transmission electron microscope (Thermo Fisher Science, Waltham, MA) at 100 kV acceleration voltage. Images were taken with a Veleta CCD camera (Olympus) operated by TIA software (Thermo Fisher Scientific). Images used for quantification were taken at 60,000x magnification. All TEM data were analyzed using Fiji imaging analysis software (Schindelin et al., 2012), RRID:SCR_002285). Linescans were made using the plot profile function in Fiji, which was also used for ssSEM and SBF-SEM analyses. Active zones (AZs) were identified by the following criteria: (1) rigid opposed pre- and post-synaptic membrane, (2) SVs clustering at the membrane, and (3) an asymmetric post synaptic density. AZs were measured as the presynaptic membrane opposing the postsynaptic density. Puncta adherentia (PA) were differentiated from AZs by lack of vesicles, a symmetric pre- and post-synaptic density, and shorter length. Vesicle distance to the AZ was calculated using a 32-bit Euclidean distance map generated from the AZ for all vesicles within 200 nm of the presynaptic membrane. Vesicles within 5 nm of the AZ were considered to be docked (Taschenberger et al., 2002; Yang et al., 2010).

### Section based immuno-electron microscopy

The samples prepared for TEM above were used for on-section immunoelectron microscopy. The MNTB region contralateral to the injected cochlear nucleus was trimmed and sectioned at 100 nm thickness using a Leica UC7 ultramicrotome. Sections were collected onto a 100-mesh nickel grid and air dried. As an antigen unmasking procedure for Durcupan resin, grids were placed section side down on water heated on a hotplate to 93-95°C for 10 minutes (Stirling and Graff, 1995). Following heat treatment, the water was slowly cooled down for 15 minutes by mixing in room temperature water. Without drying, grids were immediately placed onto drops of blocking medium for 10 minutes composed of 0.1% Tween-20, 1% bovine serum albumin, 1% normal goat serum, 1% fish skin gelatin, and 0.005% sodium azide diluted in Tris-buffered saline (TBS) buffer (pH 7.4). Grids were then incubated on a chicken anti-GFP primary antibody solution (0.5 mg/mL, Abcam Cat# ab13970, RRID:AB_300798) diluted 1:100 in the same blocking medium, and placed in a humid chamber overnight at room temperature. They were then washed and secondary antibody incubation was performed for one hour at room temperature with a 12 nm colloidal gold donkey anti-chicken antibody (Jackson ImmunoResearch Labs Cat# 703-205-155, RRID:AB_2340368) diluted at 1:30 in blocking medium. Grids were washed again on TBS and the reaction was stabilized by placing them on 1% glutaraldehyde in PBS for 5 minutes. Grids were rinsed on water and counterstained with uranyl acetate and lead citrate, and examined in a Tecnai G2 Spirit BioTwin transmission electron microscope (Thermo Fisher Scientific) at 80 kV acceleration voltage. Images were taken with a Veleta CCD camera (Olympus) operated by TIA software. Gold probe density of transduced calyces, non-transduced calyces, and principal cells was calculated by counting the number of gold particles contained within the area of each terminal or cell and dividing by that area. Raw mitochondrial intensity was calculated using Fiji by demarcating each mitochondrion within a terminal contained in a single section and calculating the mean pixel intensity across all mitochondria in the terminal.

### SBF-SEM sample processing and data analysis

Fixed brain slices of 100 µm thickness were trimmed to contain a single MNTB region and reacted with DAB and H_2_O_2_ as above. The tissue pieces were incubated in an aqueous solution of 2% osmium tetroxide and 1.5% potassium ferrocyanide for 30 minutes at 4°C in the dark. Tissue was washed with water, which was repeated between each of the following steps. Tissue was then reacted with 0.2% tannic acid in water for 30 minutes, with 1% aqueous osmium tetroxide for 30 minutes, with 1% aqueous thiocarbohydrizide for 20 minutes, and another treatment of 1% osmium tetroxide for 30 minutes all at room temperature, then placed in water overnight. The following day, tissue was stained *en bloc* with 1% phosphotungstic acid in 25% ethanol at 50°C for 30 minutes, then with 1% uranyl acetate in 25% ethanol for 30 minutes at 4°C, and Walton’s lead aspartate for 30 minutes at 60°C. At P7, staining of MNTB using this protocol was optimal for imaging. At P21, however, staining was too intense due to massive axonal myelination which readily recruits osmium. Therefore, ethanolic uranyl acetate staining was reduced to 10 minutes, and the tannic acid and second osmification steps were omitted. Infiltration and embedding were performed as above, taking extreme care in handling the brittle samples. Since SBF-SEM requires the sample to be conductive to minimize charging during imaging, the tissue was trimmed to 1 × 0.5 mm and one side was exposed using an ultramicrotome (UC7, Leica), then turned downwards to be remounted to a metal pin with conductive silver epoxy (CircuitWorks, CHEMTRONICS). The new surface was then exposed and sputter-coated with a 5 nm layer of platinum. Samples were then sectioned and imaged using 3View and Digital Micrograph (Gatan Microscopy Suite, RRID:SCR_014492) installed on a Gemini SEM300 (Carl Zeiss Microscopy LLC.) equipped with an OnPoint BSE detector (Gatan, Inc.). The detector magnification was calibrated prior to imaging within SmartSEM imaging software (Carl Zeiss Microscopy LLC.) and Digital Micrograph with a 500 nm cross line grating standard. Imaging was performed at 1.4 kV accelerating voltage, 4.45 kV extractor voltage, 30 µm aperture, working distance of ∼5 mm, 0.5 µs pixel dwell time, 6.5 nm per pixel, knife speed of 0.1–0.3 mm/sec with oscillation, and 50 or 70 nm section thickness. Images were captured at an original size of 12,000 by 12,000 pixels. The volume analyzed for whole cell reconstruction at P7 was approximately 78 × 78 × 52 µm (3.16 × 10^5^ µm^3^); the volume analyzed for P21 was approximately 78 × 78 × 79 µm, (4.81 × 10^5^ µm^3^), which was approximately the entire thickness remaining from the 100 µm tissue slice following trimming. Serial images were aligned using Digital Micrograph software and converted to TIFF format or exported as TIFFs to TrakEM2 (RRID:SCR_008954) and aligned using Scale-Invariant Feature Transform image alignment with linear feature correspondences and rigid transformation (Lowe, 2004).

For segmentation of whole terminals, images were down-sampled by a factor of four to reduce computational overhead, and only every third image was used to speed up segmentation. Aligned images were exported in TIFF format to Microscopy Image Browser (MIB; RRID:SCR_016560, (Belevich et al., 2016) for semi-automated segmentation of the calyx using the watershed brush tool with interpolation. Following segmentation, interpolation was visually verified in each section. Segmentations were saved separately and then converted to a mask for mitochondrial segmentation, allowing extract features specific to the terminal. Dark features were then thresholded in 3D and objects in 3D were morphologically filtered using the statistics function until only mitochondria remained. Segmented mitochondria were smoothed and manually proofed in each section. Volume measurements for terminals and mitochondria were calculated from meshed surfaces of the segmented data, beginning at the base of the axon, identified as the location where the area of the calyx cross section began to open into the main portion of the terminal. Axon measurements were taken from the opening of the terminal to where the axon left the volume. Only calyx terminals that were fully contained within the imaged volume were analyzed, resulting in six calyces and eight minor terminals each at P7 and P21. Scale bars presented in figures are extracted from the original EM micrographs.

For full resolution analysis of the ultrastructure of the subcompartments, one calyx at P7 (Fig. 5D far left) and two calyces at P21 (Fig. 5E far left and far right) were sampled. A P7 the calyx exists largely as a cup structure, while the P21 terminal can be divided into stalk and swelling subcompartments. We used the definition of stalk and swelling as defined by (Wimmer et al., 2006) and (Grande and Wang, 2011). An ROI measuring 6.5 × 6.5 × 7 µm was used to sample the axons, cup, and stalks, as they are not discrete structures. Swellings could be defined as discrete cytoplasmic compartments containing active zones, SVs, and typically mitochondria, having at least one thin neck connecting it to another compartment, and therefore could be sampled as distinct structures. One axon and terminal (cup) were sampled at P7. Two axons (one per calyx), four stalks (two per calyx), and 22 swellings (16 and 6 per calyx) from two calyces were sampled at P21.

For the partial volume at P7, a series of 300 sections at native XYZ resolution (6.5 nm per pixel, 50 nm thick) was used to semi-automatically segment the calyx terminal and mitochondria. A complete stack of aligned TIFF images was cropped to the size of the terminal using Fiji, then exported to MIB. Gaussian smoothing was performed on the images and the terminal was segmented using the semi-automatic graphcut segmentation function. Segmentations were proofed manually in each section.

For all reconstructions, Amira (version 6.5, Amira 3D analysis, RRID:SCR_014305) was used as a measurement tool, visualization tool, and video creation tool. Blender (version 2.79, 2.8; RRID:SCR_008606) was used to create 3D renderings of terminals.

### ssSEM sample processing and data analysis

Tissue slices were processed with the same method as for TEM above. Once embedded, samples were trimmed to a roughly 2 × 3 mm rectangle, sectioned at 50 nm using the ATUMtome (RMC Boeckeler, Tuscon, AZ), and collected on a reel of Kapton tape. Tape containing sections was attached to a 4-inch silicon wafer in rows using double sided carbon tape (EMS, Cat# 77817-50) and sections were post-stained with uranyl acetate and lead citrate. Wafers were coated with a 5 nm layer of carbon before imaging to limit charging (Leica, Wetzlar, Germany, ACE600). Image acquisition was performed semi-automatically using a Merlin VP Compact SEM equipped with Atlas 5 AT software (Carl Zeiss Microscopy LLC., Oberkochen, Germany). Overview images of sections were acquired and registered using SE2 signal at 2 µm per pixel. Medium magnification images of the general region of interest were acquired to identify potential synaptic terminals at 30 nm per pixel using a BSE detector. High magnification images of targets were captured using an Inlens Duo detector at an accelerating voltage of 1.9 kV, 4.45 kV extractor voltage, 60 µm aperture, ∼6.8 mm working distance, ∼12.8 µs pixel dwell time, and 5 nm per pixel. The detector was calibrated prior to imaging using a 500 nm cross line grating replica. Images were batch exported as TIFF files and aligned automatically using the TrakEM2 plugin using Scale-Invariant Feature Transform image alignment with linear feature correspondences and rigid transformation. Microscopy Image Browser (MIB) was used to semi-automatically segment calyx of Held terminals using the watershed brush tool. Amira was used for analysis of the 3D vesicle pool, vesicle docking, and extraction of AZ surface areas; as a visualization and video creation tool. All vesicles within a 200 nm distance from AZ were manually selected from the image stack, and a custom script was used to calculate the nearest distance from each vesicle to the surface of the AZ. AZ surface areas were extracted from the presynaptic side of the triangular mesh formed following reconstruction. Estimates of the total number of AZs and PA were quantified using MIB.

### Controls for EM analysis

Internal controls were taken from the non-transfected MNTB ipsilateral to the injection site, or, if expression was not ubiquitous in the contralateral MNTB, individual cells identified as being negative for EGFP expression and by the lack of electron dense mitochondria (compared to nearby infected cells) were used. For post-embedded samples, lack of gold particles labeling EGFP was also used to confirm a terminal was uninfected. We did not observe ipsilateral expression of EGFP or mito-APEX2 in any processed tissue by fluorescence, DAB reaction product, or EM dense staining at the mitochondrion. For linescan analyses, principal neuron mitochondria were used as a staining control. For the immuno-EM study, antibody specificity was examined by omitting primary antibody from the diluent solution during the staining protocol.

### Experimental Design and Statistical analysis

For the electrophysiological data analysis, individual neurons were considered independent samples. For immuno-TEM analysis, different cell compartments from one MNTB region (i.e. calyx terminals or principal cell bodies) were considered independent samples. For thin-section TEM analysis, AZs were considered independent samples 120 total AZs, (3 animals, 40 AZs each). Data were analyzed in MATLAB (version 9.4, RRID:SCR_001622), GraphPad Prism (RRID:SCR_002798), R Project for Statistical Computing (version 3.5.2, RRID:SCR_001905), and JASP (version 0.9.2.0, RRID:SCR_015823). Data distributions were tested for Gaussianity using the Shapiro-Wilk test. To compare two groups, a two-tailed unpaired Student’s t test with Welch’s correction (normal distribution) or two-tailed Mann-Whitney *U* test (non-normal distribution) were used. Immuno-TEM data groups were compared using Mood’s median test. Linear regressions of total mitochondria volume to terminal volume were fit through the origin of the axes. For mitochondria volume ratio measurements of calyx terminals, a two-way repeated measures ANOVA was conducted to compare the effects of age and compartment on volume ratio. Mitochondria diameter was compared using one-way ANOVA. P-values from multiple comparisons were adjusted using the Bonferroni method. Data are reported as mean ± standard error of the mean, unless otherwise stated. For interpretation of all data, a *P*-value of 0.05 was deemed significant. *P*-values in figures are denoted \**P*<0.05; \*\**P*<0.01; \*\*\**P*<0.001; \*\*\*\**P*<0.0001. Since non-significant *P*-values do not provide evidence for the lack of effect or to draw conclusions about the probability of the null hypothesis (H0) over the alternative hypothesis (H1), we additionally calculated the Bayes Factor using a Cauchy distribution with scale factor 1 as the prior distribution implemented as described previously (Rouder et al., 2009). The Bayes Factor is reported as BF_10_ for immuno-TEM indicating the likelihood of the data under H1 (treatment has an effect) relative to H0 (treatment has no effect), where values larger than 1 are in favor of H1. For thin-section-TEM and electrophysiology, BF_01_ is reported, indicating the likelihood of the data under H0 (treatment has no effect) relative to H1 (treatment has an effect), where values larger than 1 are in favor of H0. Effect sizes were calculated using the MES toolbox in MATLAB (Hentschke and Stuttgen, 2011) and are reported as Cohen’s U_1_ (proportion of non-overlap of the two distributions) for two-sample comparisons and eta-squared (η^2^, the proportion of the total variation explained by the variation between groups) for multiple-sample comparisons. Confidence intervals of effect sizes were calculated using bootstrapping with 10,000 repetitions. No statistical test was performed to pre-determine sample sizes. Exact *P*-values, test statistics, Bayes Factors, and effect sizes for all statistical comparisons are summarized in Table 1.

### Data and Software Accessibility

All experimental data, movies, 3D reconstructions and custom-written software central to the conclusion of this study are available at http://dx.doi.org/10.17632/v88r5t5myz under the terms of the Creative Commons Attribution 4.0 License (CC BY 4.0).

## Results

### HdAd EGFP mito-APEX2 transduction leads to efficient labeling of mitochondria at the calyx of Held

To specifically label mitochondria in the calyx we created an HdAd which expresses mito-APEX2 and EGFP under the control of the hsyn promoter (HdAd mito-APEX2) (Fig. 1A1). This dual expression allowed for identification of transduced calyces with both electron and fluorescence microscopy. Stereotactic injection of HdAd mito-APEX2 into the anteroventral cochlear nucleus (aVCN) of C57Bl/6J mice was performed at postnatal day 1 (P1) as previously described (Chen et al., 2013) and experiments were carried out on P7 or P21 calyces representing the immature and mature calyx, respectively (Fig. 1A2) (Borst and Soria van Hoeve, 2012). HdAd injection into the aVCN resulted in the transduction of calyces which were identified by EGFP expression (Fig. 1B). We confirmed that these EGFP positive terminals contained APEX2-tagged mitochondria by carrying out DAB staining which resulted in dark reaction products in resin-embedded tissue under brightfield illumination. As expected, only the MNTB contralateral to the injection site contained DAB reaction products (Fig. 1C, D). At high magnification, individual calyx terminals were visible as areas of dark deposition surrounding the somata of principal cells (Fig. 1E, F). To confirm that these electron dense precipitates were sufficient to identify mitochondria in the transduced calyx terminals, we imaged ultrathin sections using TEM. We found that presynaptic calyx terminals in both P7 and P21 samples contained electron-dense mitochondria with DAB precipitates accumulated in the mitochondria matrix while the postsynaptic cell bodies did not (Fig. 1G-I). Additionally, labeled calyx mitochondria were easily detectable and distinguishable from unlabeled calyx mitochondria (Fig. 1J, K). This indicates that expression of mito-APEX2 using HdAd is sufficient to label mitochondria for EM applications.

**Figure 1.**
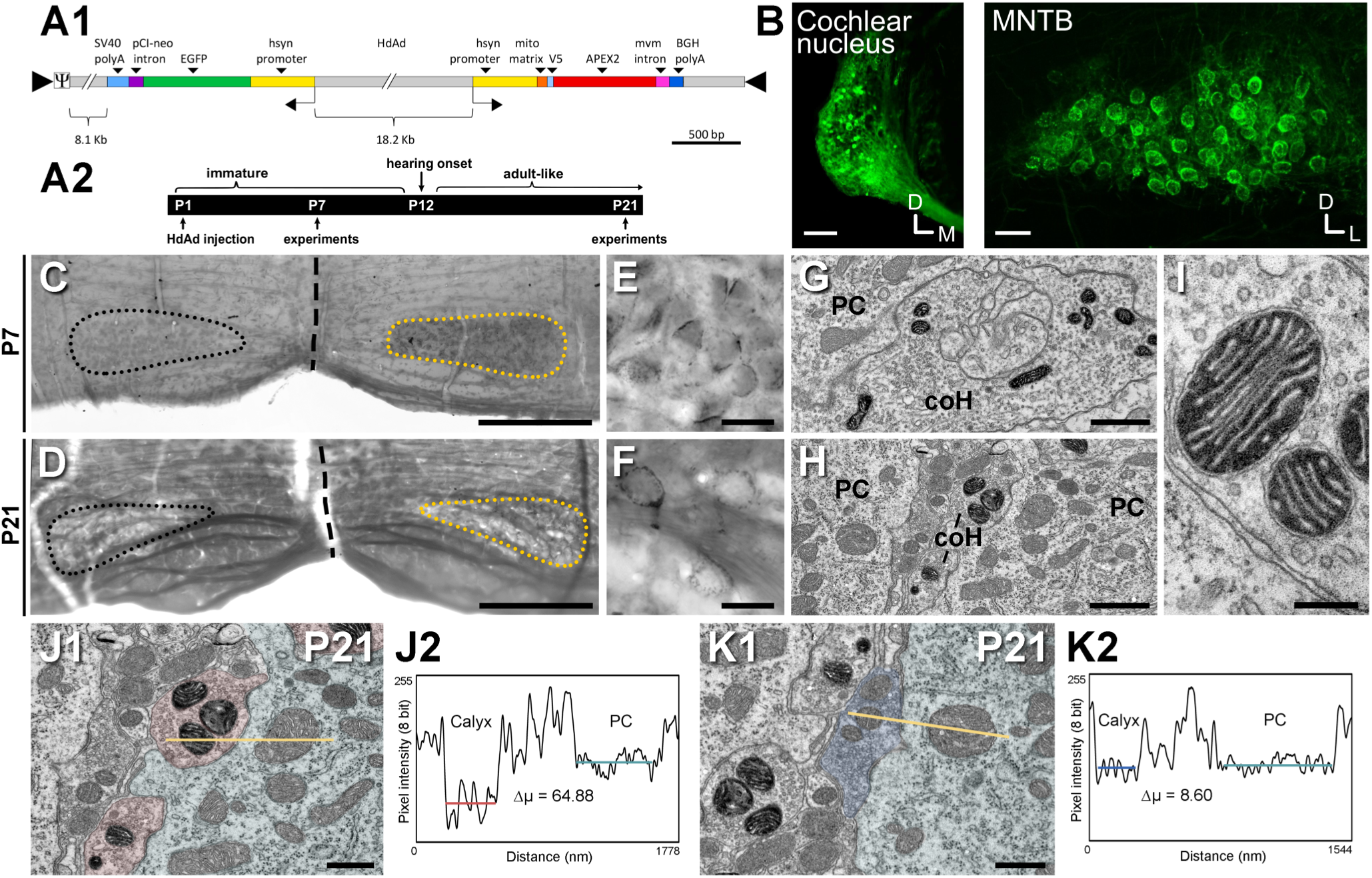
Expression of EGFP-mito-APEX2 at the calyx of Held presynaptic terminal. ***A1***, Schematic of the DNA construct encoding EGFP (green) and mitochondrial matrix-targeted APEX2 (red). ***A2***, Developmental (top) and experimental (bottom) timeline of the calyx around the onset of hearing (P12). Mito-APEX2 expressing HdAds were injected at P1 and experiments performed at either P7 or P21. ***B***, Expression of EGFP following HdAd injection into the anteroventral cochlear nucleus resulted in green fluorescence in both the cochlear nucleus (left) and the calyx of Held terminals in the contralateral MNTB (right). Images were acquired at P21. ***C***, ***D***, Following DAB reaction, mito-APEX2 labeling of the contralateral MNTB was visible in embedded P7 (***C***) and P21 (***D***) tissue slices under brightfield microscope. Dashed line represents the slice midline, dotted lines demarcate the MNTB region. Yellow dotted line represents the transduced MNTB region contralateral to the injection site. ***E***, ***F***, High magnification brightfield images of transduced calyx terminals from ***C*** and ***D***. Mito-APEX2 expressing calyces where visible by the dark area around the MNTB principal cell soma. ***G***, ***H***, Transmission electron micrographs at P7 (***G***) and P21 (***H***) of transduced calyx of Held terminals showing electron dense mitochondria in juxtaposition to mitochondria in uninfected MNTB principal cells. ***I***, High magnification image of labeled mitochondria in a transfected calyx terminal showing labeling is restricted to the matrix of the mitochondrion, while the intermembrane space remains unstained. ***J1***, Transduced terminals at P21 (red) contain darker mitochondria (having higher electron density) compared to mitochondria of the opposing principal cell (aqua). ***K1***, Non-transduced terminal (blue) mitochondria compared to the principal cell. ***J2***, ***K2***, Intensity line profiles of mitochondrial intensity in transduced and non-transduced terminals, Δµ = difference in mean pixel intensity between calyx and PC mitochondria. coH, calyx of Held; PC, principal cell. Scale bars: ***B***, 200 µm, 50 µm; ***C***, ***D***, 300 µm; ***E***, ***F***, 25 µm; ***G***, ***H***, 1 µm, ***I***, 200 nm. ***J***, ***K***, 500 nm.

### EGFP and mito-APEX2 dual expression does not alter presynaptic ultrastructure

To confirm and characterize the dual expression of EGFP and mito-APEX2 we performed post-embedding immunogold labeling of EGFP on ultrathin sections from P21 calyces labeled for mito-APEX2 (Fig. 2). The cytoplasm of all calyx terminals containing electron dense mitochondria also contained gold particle labeling of EGFP (Fig. 2A, B). To quantify the expression of EGFP and mito-APEX2, we analyzed gold particle density in calyces containing labeled mitochondria compared to non-transduced calyces and principal cells (Fig. 2C). Immunogold particle density was highest in terminals containing mito-APEX2 labeled mitochondria (terminal control = 2.70 ± 0.37 particles/µm^2^, principal cell control = 3.55 ± 0.34 particles/µm^2^ vs. EGFP-mito-APEX2 = 14.48 ± 1.10 particles/µm^2^, *P* = 0.00015; Fig. 2C, Table 1). Although mito-APEX2 was constrained to mitochondria, we tested if its expression affected presynaptic ultrastructure. Using ultrathin sections from P21 animals, we analyzed active zone (AZ) length, SV distribution, and docked SV number (within 5 nm of the presynaptic membrane) (Taschenberger et al., 2002) for both mito-APEX2 expressing calyces and their respective in-slice control. We observed no significant difference in any measured parameter between control and mito-APEX2 expressing terminals (Fig. 2D-F and Table 1).

**Figure 2.**
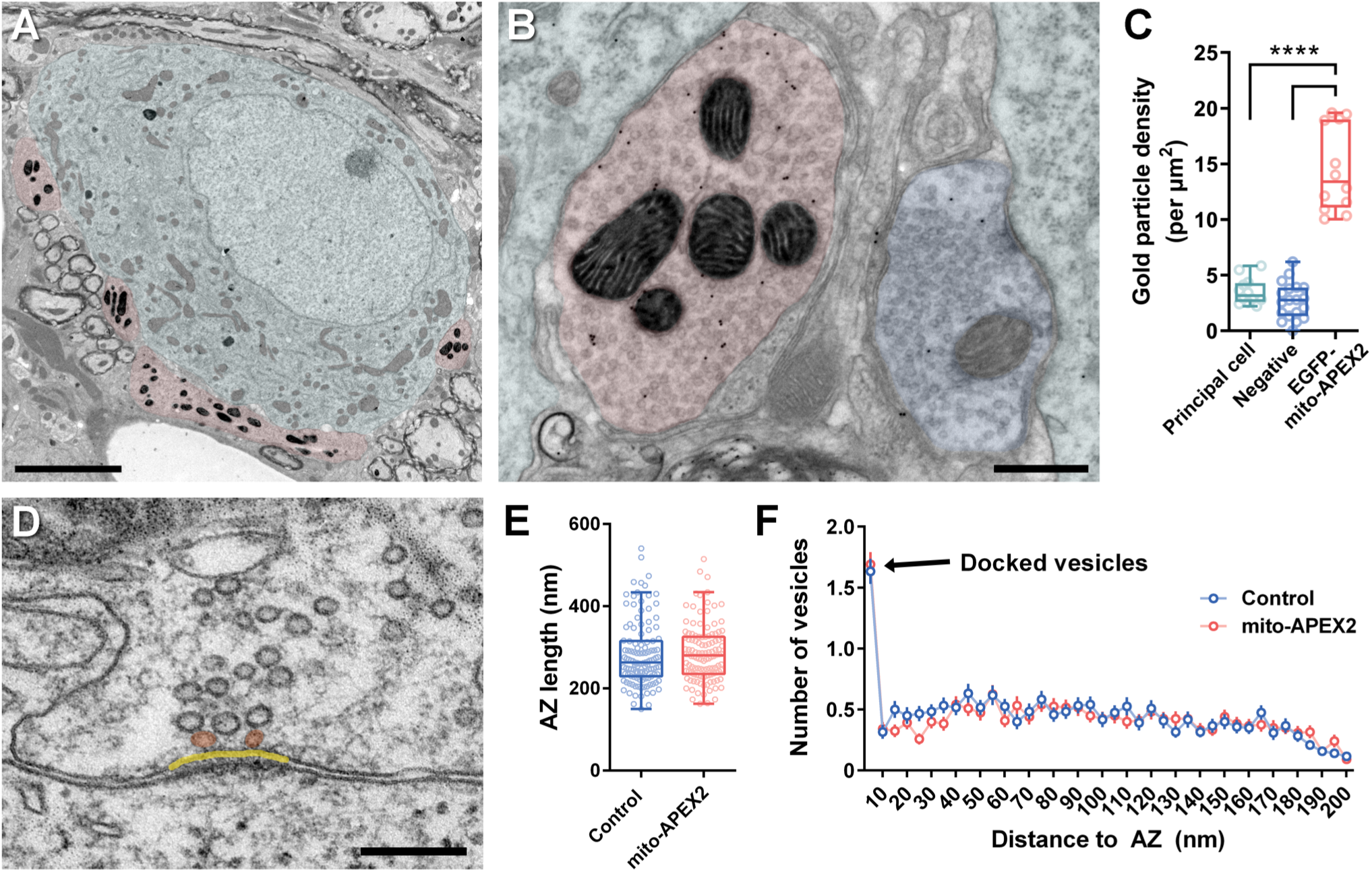
Double labeling of EGFP and mito-APEX2 using a combination of pre-embedding DAB-reaction labeling and post-embedding immunogold labeling. Expression of EGFP and mito-APEX2 has no effect on ultrastructure. ***A***, Transduced calyx of Held terminal (red) at P21 with electron-dense mitochondria makes synaptic contact with a single principal cell (aqua). ***B***, Comparison of a transduced calyx terminal expressing mito-APEX2 (red) versus a non-transduced control terminal (blue), both contacting separate principal cells (aqua). 12 nm gold particles labeling EGFP are visible within the presynaptic terminal of transduced cells. ***C***, Gold particle density is significantly higher in calyx terminals expressing EGFP and mito-APEX2 compared to non-transduced negative terminals and principal cells. ***D***, Two docked vesicles (orange) convene at a synaptic release site. Active zone is demarcated in yellow. Vesicles less than 5 nm from the active zone were consider docked. ***E***, Active zone length is unchanged between control and mito-APEX2 expressing terminals. A total of 120 AZs from control and mito-APEX2 expressing terminals of three different animals (P21) were analyzed. ***F***, Single-section distribution of SVs up to 200 nm from the AZ. Box plot whiskers extend to the minimum/maximum within 1.5 interquartile range, open circles indicate individual observations. Scale bars: ***A***, 5 µm; ***B***, 500 nm, ***D***, 200 nm, 200 nm

### Mito-APEX2 expression does not alter synaptic transmission

HdAd mito-APEX2 lead to efficient mitochondria labeling in the calyx and ability to visualize presynaptic ultrastructure. However, it has been shown that extreme levels of APEX expression can impact mitochondria structure (Martell et al., 2017). Since proper mitochondria function is integral for synaptic transmission and plasticity (Billups and Forsythe, 2002; Cserep et al., 2018; Lee et al., 2018), we tested if mito-APEX2 expression impacts synaptic transmission. First, we performed afferent fiber stimulation of transduced calyces at P7 and recorded whole-cell AMPAR currents of MNTB principal cells using 50 Hz stimulation. We did not find evidence for altered synaptic transmission compared to age-matched WT control (Fig. 3A, Table 1). Finally, we tested if chronic expression of HdAd mito-APEX2 impacted synaptic transmission and repeated the experiments with P21 animals. Using afferent fiber stimulation at 300 Hz, we found that chronic mito-APEX2 expression did not alter synaptic transmission (Fig. 3B, Table 1). These data combined with the ultrastructural measurements indicate that chronic expression of mito-APEX2 does not alter presynaptic ultrastructure or synaptic transmission.

**Figure 3.**
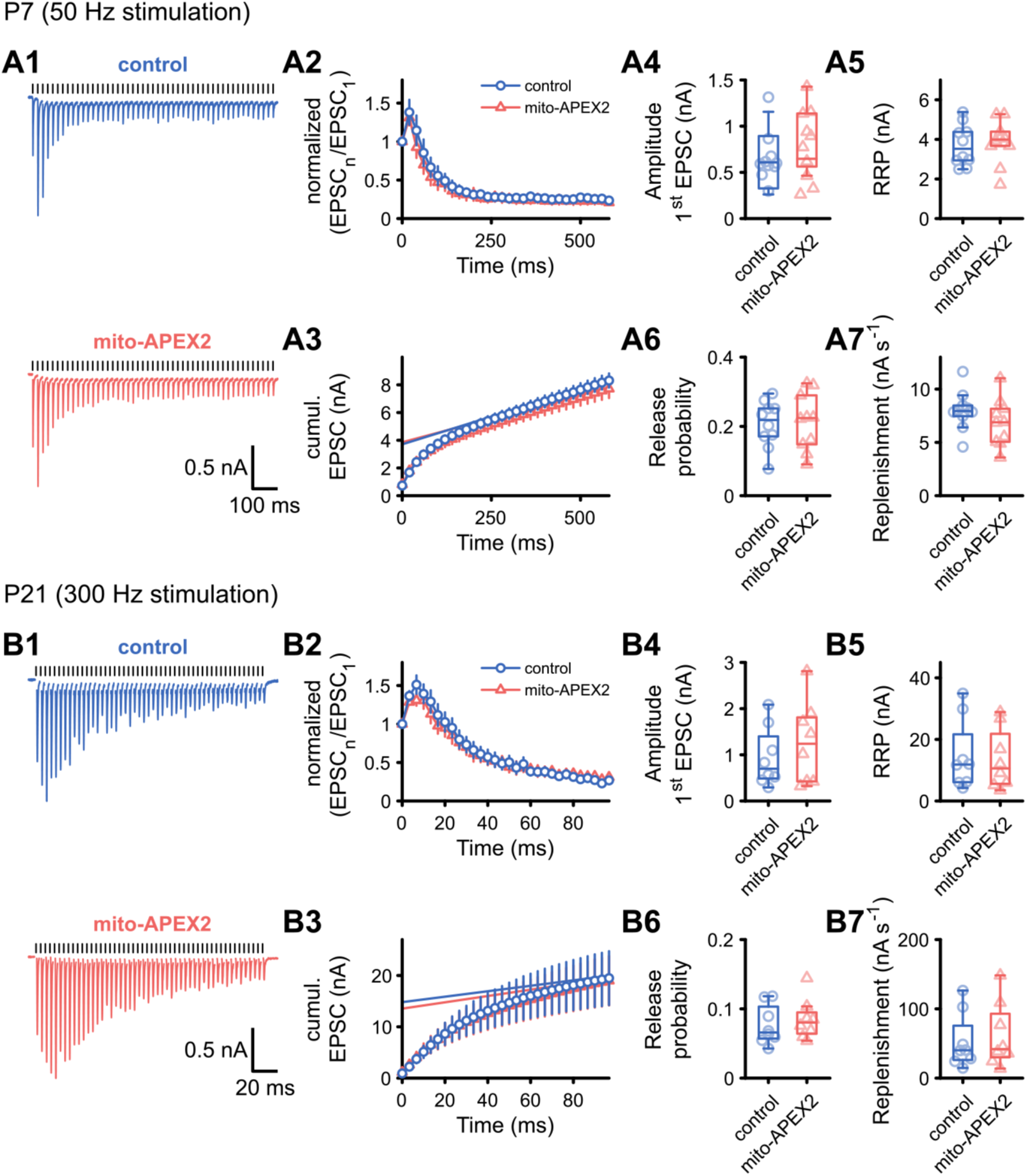
Expression of mito-APEX2 does not affect synaptic transmission at the calyx of Held-MNTB synapse. Synaptic transmission at the calyx of Held after expression of mito-APEX2 (red) was investigated at P7 (***A***) and P21 (***B***) and compared to age-matched wildtype control (blue). ***A1***, Representative traces of evoked EPSCs during afferent fiber stimulation (50 Hz) at P7 for control and mito-APEX2-transduced calyces (injected at P1). Stimulus artifacts are blanked. Vertical black lines above the traces indicate time points of fiber stimulation. ***A2***, Averaged EPSC amplitudes normalized to the first EPSC showed comparable short-term plasticity between control and mito-APEX2-transduced calyces. ***A3***, Cumulative EPSC amplitudes to determine the readily releasable pool (RRP) by back-extrapolating the linear fit of the last 10 data points; the y-intercept is an estimate of the RRP size. ***A4-A7***, No differences between control and mito-APEX2-transduced calyces were detected regarding EPSC amplitude (***A4***), RRP size (***A5***), release probability (***A6***), or replenishment rate (***A7***). ***B,*** Same layout as in ***A***. Afferent fiber stimulation was performed at 300 Hz. Following chronic expression (20 days) of mito-APEX2, the calyx of Held-MNTB synapse showed normal synaptic transmission and no change in EPSC amplitude (***B4***), RRP (***B5***) release probability (***B6***), or replenishment rate (***B7***). Box plot whiskers extend to the minimum/maximum within 1.5 interquartile range, open circles indicate individual cells.

### Large scale volumetric analysis of calyces and mitochondria with SBF-SEM

Due to the large size of the calyx, volumetric analysis at fine-scale EM resolution of mitochondria has been restricted to partial reconstructions of single calyces using TEM (Rowland et al., 2000; Wimmer et al., 2006), ssSEM (Horstmann et al., 2012), or electron tomography (Perkins et al., 2010). Furthermore, ultrathin section-based imaging platforms are constrained by alignment issues of single sections which hampers large scale volumetric analysis (Lichtman and Denk, 2011). In contrast, SBF-SEM requires minimal image alignment and is therefore ideal to efficiently image large EM volumes. Although SBF-SEM has been successfully used for calyx reconstructions (Holcomb et al., 2013; Xiao et al., 2013) it requires liberal application of heavy metals, which may prevent detection of mito-APEX2 labeling. To test this, we used the 3View with OnPoint BSE detector and optimized the imaging conditions (see Methods) to confirm that mito-APEX2 labeled mitochondria could be distinguished in both P7 and P21 (Fig. 4). We found calyx terminals containing mito-APEX2 positive mitochondria were clearly identifiable in both P7 and P21 samples as the electron density of labeled mitochondria in the calyx terminal was considerably higher than that of unlabeled mitochondria in the principal cell (Fig. 4E, F). Although we could visualize SVs at this resolution, identification of AZs (using the postsynaptic density as a reference) and PAs was difficult due to the lower signal-to-noise ratio of images acquired with short dwell times (Fig 4B, D).

**Figure 4.**
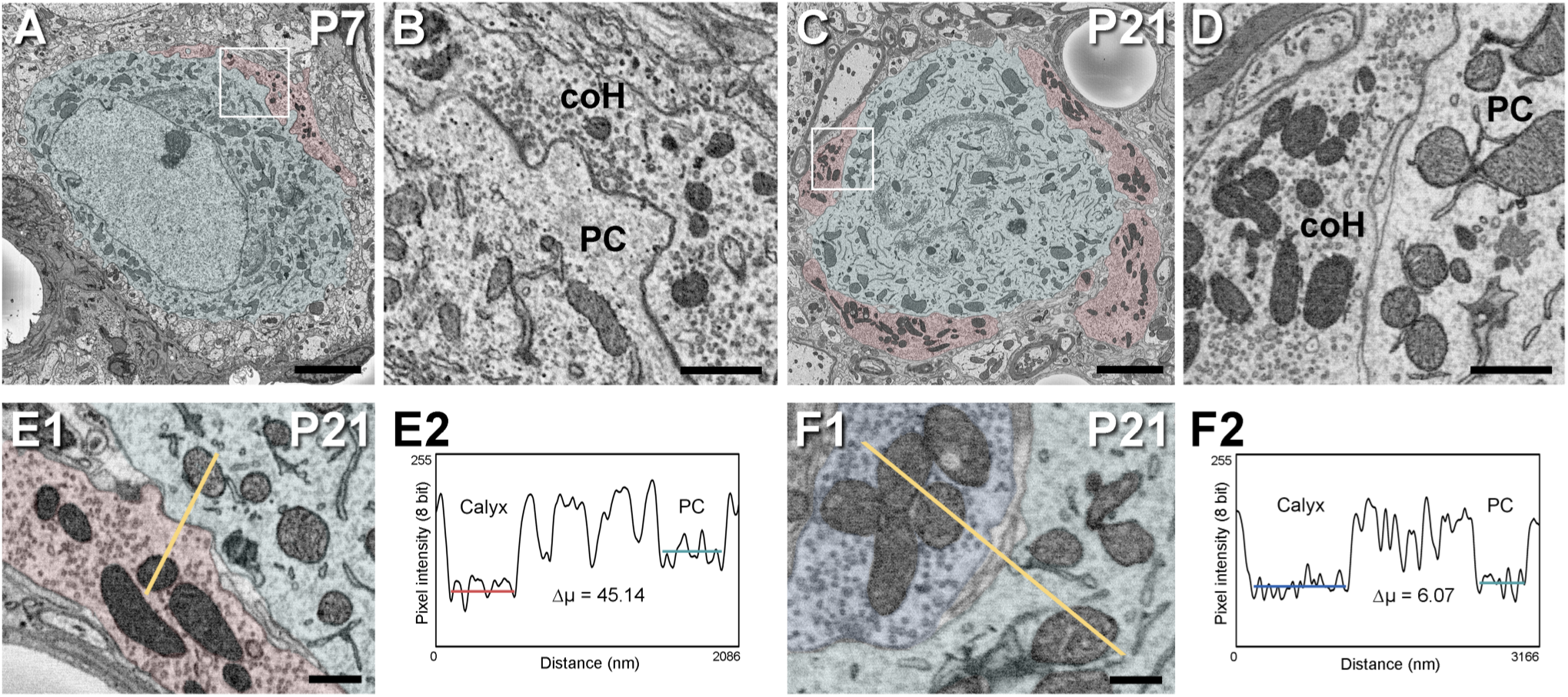
Mito-APEX2 labeling visualized by SBF-SEM at P7 (***A***, ***B***) and P21 (***C***, ***D***, ***E1***) allows for identification of transduced calyx of Held terminals. ***A***, ***C***, Block face images show the calyx of Held (red) contacting to an MNTB principal cell (aqua). ***B***, ***D***, High magnification image of the white squared area in ***A*** and ***C***, respectively, showing electron dense mitochondria in the calyx of Held terminal. ***E1***, Transduced terminals contain darker mitochondria compared to mitochondria of the opposing principal cell. ***F1***, Non-transduced terminal mitochondria compared to the principal cell mitochondria. ***E2***, ***F2***, Intensity line profiles of mitochondrial intensity in transduced and non-transduced terminals, Δµ = difference in mean pixel intensity between calyx and PC mitochondria respectively. coH, calyx of Held; PC, principal cell. Scale bars: ***A***, ***C***, 5 µm; ***B***, ***D***, 1 µm; ***E1***, ***F1***, 500 nm

Subsequently, we acquired and analyzed volumes of 78 × 78 × 52 µm (P7, Fig. 5A) and 78 × 78 × 79 µm (P21, Fig. 5B) at an XY pixel size of 6.5 nm and section thickness of 70 nm. We found that a pixel dwell time of 0.5 µs provided sufficient image quality for tracing membranes while reducing imaging time, which allowed us to collect the full volume in ∼30 hours per sample. We segmented individual positive calyx terminals that were within the bounds of the datasets and carried out whole terminal reconstructions. To increase segmentation throughput and minimize dataset size, images were down-sampled and every third image was used. In the acquired image volume, all six calyx terminals containing mito-APEX2 positive mitochondria plus eight labeled minor inputs were segmented at P7 in ∼40 working hours. Due to the complex calyx structure in P21 animals, segmentation of six terminals and eight labeled minor inputs required ∼100 working hours. In those volume data sets, the reconstructed terminals at P7 and P21 showed clear morphological differences as previously described (Satzler et al., 2002; Taschenberger et al., 2002; Wimmer et al., 2006; Grande and Wang, 2011; Horstmann et al., 2012; Holcomb et al., 2013) (Fig. 5A, B, and D, E, Movie 1 and 2). At P7, several filopodial extensions branched off from the cup-like terminal (Fig. 5D). These cups were thin and sheet-like, while terminals at P21 were composed of mainly thick, bulbous compartments without filopodial extensions (Fig. 5E). We measured the volume of individual calyx terminals containing mito-APEX2 positive mitochondria and found that calyx volumes at P21 were larger than at P7 (1307 ± 124 µm^3^ vs. 841.2 ± 136.1 µm^3^, *P* = 0.03; Table 2, Fig. 5C). The terminals at P7 had volumes between 9 and 1171 µm^3^ and 5 of the 6 MNTB principal neurons were contacted by a large calyx presynaptic terminal (>700 µm^3^) with minor terminals, while one principal neuron had a dominant, but smaller calyx terminal of ∼250 µm^3^ and multiple minor terminals (<100 µm^3^) (Fig. 5A, D, yellow terminals, Movie 1). Consistent with our data, principal cells with small dominant calyx and multiple minor competing terminals are rare at P7, comprising only roughly 20% of cells (Holcomb et al., 2013).

**Figure 5.**
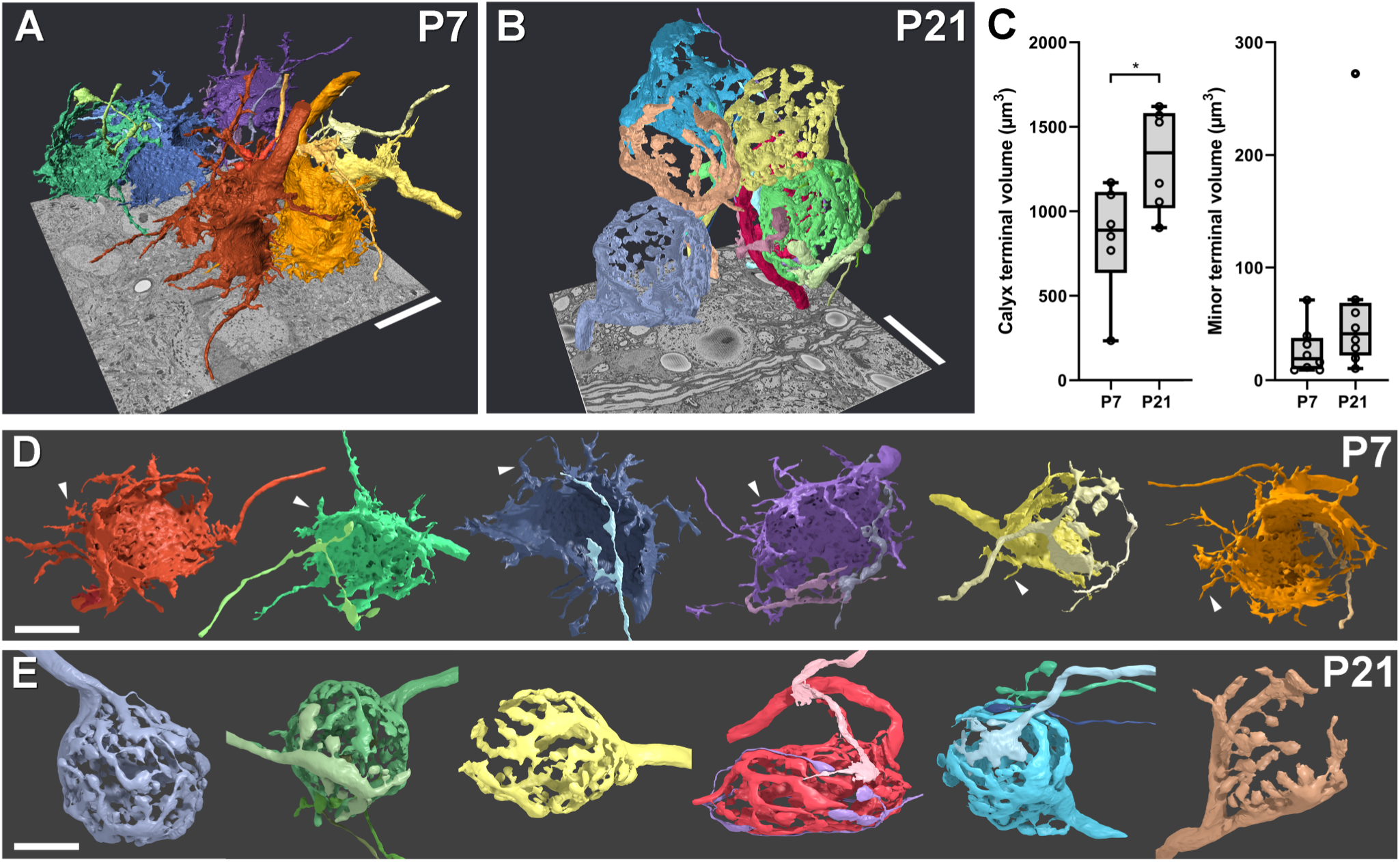
Large volume reconstructions of mito-APEX2-positive calyx of Held terminals using SBF-SEM at P7 (***A***, ***D***) and P21 (***B***, ***E***). ***A***, Volumetric positions of six transduced calyx terminals and eight minor inputs contacting six different MNTB principal cells (not shown), reconstructed from a down-sampled volume of 78 × 78 × 52 µm at P7. ***B***, Volumetric positions of six transduced terminals with eight minor inputs contacting six principal cells (not shown) reconstructed from a down-sampled volume of 78 × 78 × 79 µm at P21. ***C***, Volumes of calyx terminal (left) and minor terminal (right) at P7 and P21. ***D***, Individual renderings of mito-APEX2-positive calyx terminals together with 1-3 minor labeled terminals contacting each principal cell (not shown). Several filopodia extend from the cup-like calyx terminals (examples are pointed with white arrowheads). ***E***, Individual renderings of mito-APEX2 positive calyx terminal inputs contacting principal cells (not shown). Thick digitated terminals at P21 had few filopodial extensions, but 3 out of 6 principal cells had minor labeled inputs (migratory calycigenic axons; second, fourth, and fifth from left). Box plot whiskers extend to the minimum/maximum within 1.5 interquartile range, open circles indicate individual terminals. Scale bars: ***A***, ***B***, 20 µm; ***D***, ***E***, 10 µm.

At P21, terminals ranged from 10 to 1620 µm^3^. We found that all (6/6) principal cells at this age were contacted by a single dominant calyx similar to previous reports (Holcomb et al., 2013) and exhibited a markedly different structure with variable degrees of fenestration as described previously (Grande and Wang, 2011; Wang et al., 2015); Fig. 5E, Movie 2). In addition, half of those principal cells (3/6) had mito-APEX2 labeled minor terminals (Fig. 5E, Table 2), with morphologies similar to “migratory calycigenic axons” in the young opossum (Morest, 1968) and calyceal collaterals in rodents (Rodriguez-Contreras et al., 2006). Notably, one principal cell had a moderately sized terminal (272 µm^3^) in addition to minor terminals (Fig 5B, E, green terminals). Therefore, we conclude that even though principal cells are innervated by a dominant calyx, minor terminal still exist in the functionally mature stage of development, which might be comprised by calyceal collaterals. Finally, in addition to the labeled minor terminals, numerous small unlabeled terminals contacted the same principal neurons at both ages but were not segmented or analyzed. These terminals may compose additional excitatory or inhibitory inputs to the MNTB principal cell that do not originate from the cochlear nucleus (Hamann et al., 2003; Awatramani et al., 2004; Albrecht et al., 2014).

### Mitochondrial volume ratios are higher in the mature calyx and axon than the immature calyx

Since the high-contrast mitochondria could be easily extracted using thresholding techniques, we carried out high resolution whole terminal mitochondria reconstructions at P7 and P21 using the same volume data sets acquired with SBF-SEM. After down-sampling the volume data and using every third image, whole terminal mitochondrial volume reconstructions could be completed and proofed in several hours per terminal (Fig. 6A-D, Table 2, Movies 3 and 4). Since at both developmental stages, principal cells had labeled minor terminals, we separately analyzed mitochondrial volume in the calyx terminal and the minor terminals. Mitochondrial volumes scaled linearly with terminal volume at both developmental time points. However, we found that the scaling of mitochondria to calyx volume is steeper at P21 than at P7 (Slope = 0.2022, R^2^ = 0.95 vs. 0.0737, R^2^ = 0.82; Fig 6E). Comparison of the mitochondria to volume ratio of the terminal revealed that 20% of the P21 calyx volume is composed of mitochondria, while at P7 mitochondria only take up 7.8% of the terminal—a ∼2.6-fold increase throughout development (*P <* 0.0001; Fig. 6E, Table 2). Mitochondrial volumes in the minor terminals at both P7 and P21 showed a similar linear scaling of mitochondrial volumes (Slope = 0.1883, R^2^ = 0.99 vs. 0.1066, R^2^ = 0.95) and revealed a ∼40% increase in the mitochondrial volumes at P21 (*P* = 0.0245; Fig. 6F).

**Figure 6.**
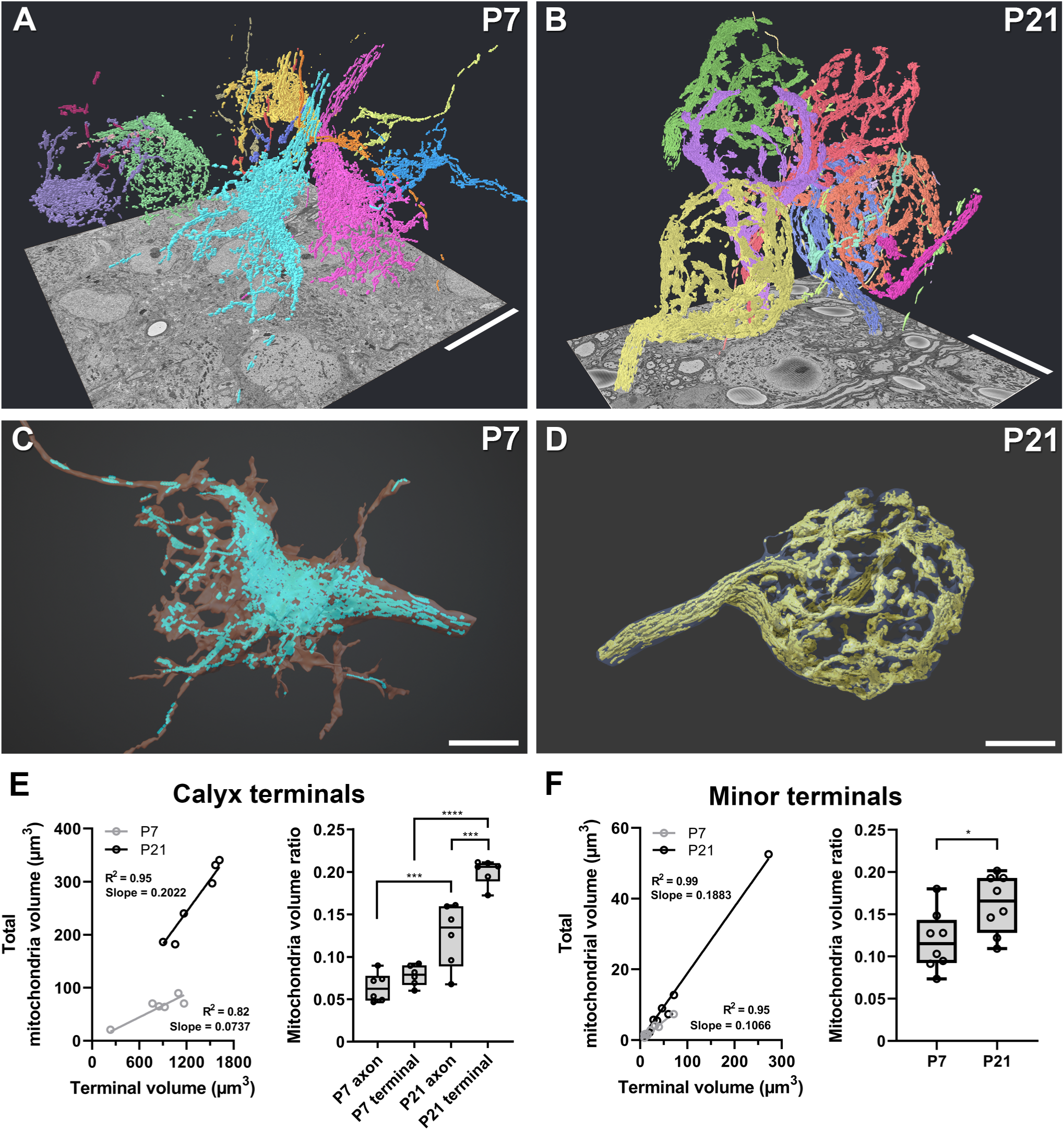
Volumetric reconstructions of mitochondria from the same whole terminals and axons shown in Figure 5 using SBF-SEM at P7 and P21. A, B, Volumetric positions of mito-APEX2-labeled mitochondria of the six calyx of Held terminals at P7 (A) and P21 (B). C, D, individual renderings of mito-APEX2-labeled mitochondria with the membrane surface view of the whole terminals in transparent gray from one terminal at P7 (C) and P21 (D). E, Mitochondrial volume scales linearly with the calyx terminal volume at both developmental stages. Mitochondrial volume ratio is not different between axons and terminals at P7, but volume ratio does increase in axons and terminals across developmental stages. Ratios are also higher in P21 terminals compared to the P21 axons. F. Mitochondrial volume scales linearly with terminal volume in minor terminals at both developmental stages. Mitochondria volume ratios are slightly higher in P21 minor terminals compared to P7. Box plot whiskers extend to the minimum/maximum within 1.5 interquartile range, open circles indicate individual axons or terminals. Scale bars: A, B, 20 µm; C, D 10 µm

During maturation, the calyx transforms into a high-fidelity synapse that can sustain action potential (AP) firing upwards to a kilohertz (Baydyuk et al., 2016). Since AP firing is an energetically expensive process (Harris et al., 2012), this suggested that mitochondria levels are increased in the axon also. To determine if mitochondria volumes are increased in the axon to support the increased firing rates during development, we analyzed the mitochondrial volumes in the calyx axon. We found that at P7 mitochondria composed 6.4% of the axon volume, while at P21 the mitochondria volume was increased 2-fold, comprising 12.5% of axon volume (*P* = 0.0003). When comparing the relative mitochondrial volumes in the calyx terminal vs. axon, we found that the calyx had significantly higher relative mitochondria volumes than the axon at P21 (*P* = 0.0009), while there was no difference between the P7 terminal and axon (*P* = 0.7116; Fig. 6E, Table 2).

Taken together, our results revealed that mitochondrial volumes increase in both the calyx and its axon during maturation. Furthermore, mitochondrial volumes in the mature calyx are selectively increased compared to its respective axon, resulting in an asymmetry not found in the immature calyx. Therefore, we conclude that increases in mitochondrial volumes support the energetic demands required for high fidelity synaptic transmission.

### Stalks and swellings of the mature calyx have similar relative mitochondrial volumes

In the mature calyx, Ca^2+^ transients are higher in swellings compared to stalks, and stalks have higher *Pr* than swellings (Fekete et al., 2019). Since mitochondria can act as Ca^2+^ buffers, this suggests mitochondria volumes may be lower in swellings compared to stalks. In addition, swellings have more loosely coupled SVs than stalks which are proposed to be in a less energetically favorable configuration (Eggermann et al., 2012). Therefore potential differences in coupling may impact mitochondrial volumes between the two subcompartments. To determine mitochondrial volumes in these subcompartments we segmented small subvolumes at full resolution and reconstructed mitochondria in 4 stalks and 22 swellings from two calyces (P21, far left and far right calyces in Fig. 5E; 7C-E). Subsequent analysis revealed that stalks and swellings contain similar relative volumes of mitochondria (stalks = 17.5% vs. swellings = 21.4%, Fig. 7D, E, Table 2).

**Figure 7.**
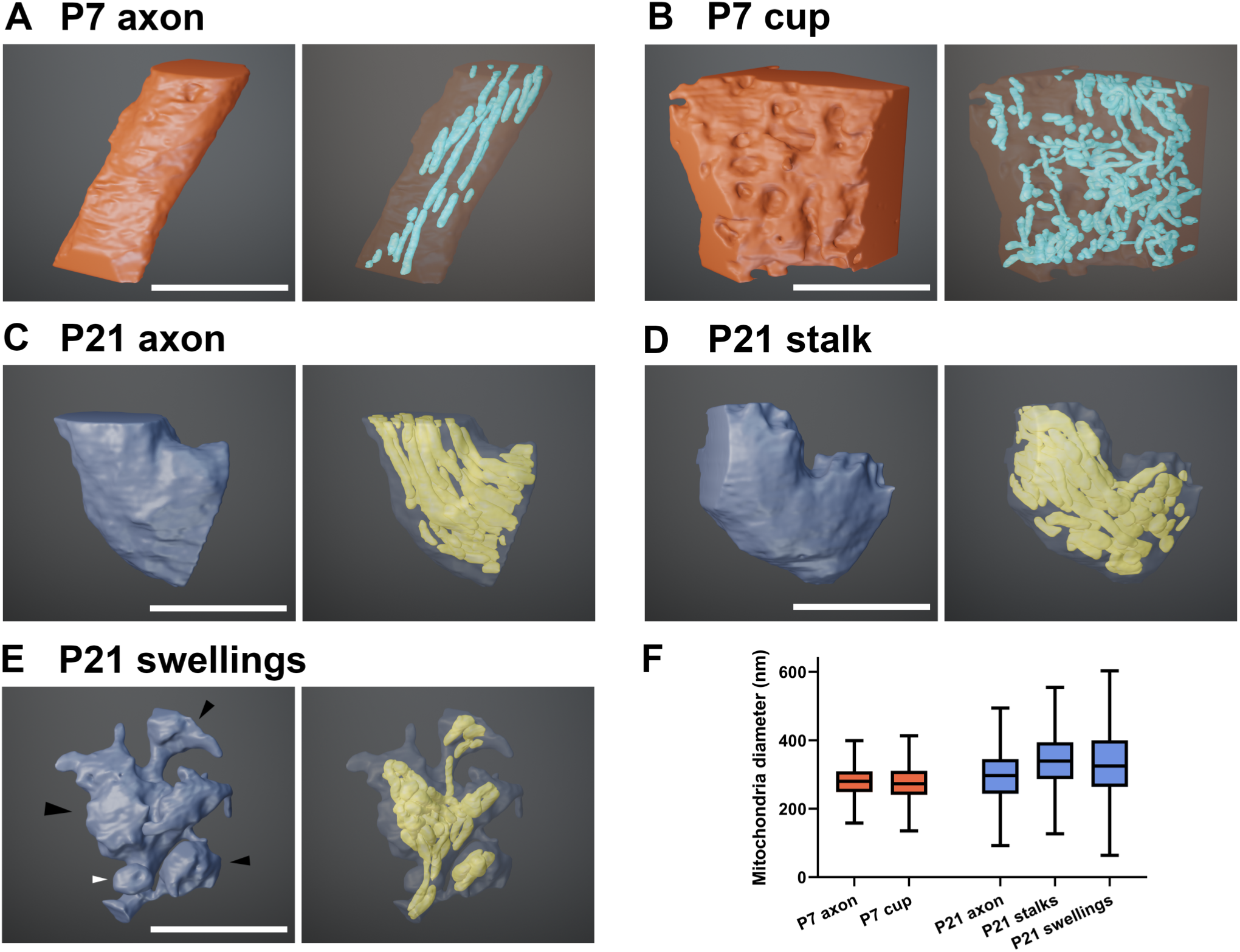
Full resolution reconstructions of subcompartments and mitochondria of calyx of Held terminals at P7 and P21 using SBF-SEM. ***A-E***, Reconstruction of a portion of P7 calyx of Held axon (***A***) and terminal (cup) (***B***); and example reconstructions of a portion of P21 axon (***C***), stalk (***D***), and swellings (***E***). Left images show the membrane surface views of the subcompartments and right images show the mitochondria inside. Black arrowheads (***E***) point to several individual swellings of different sizes. White arrowhead indicates a small swelling not containing mitochondria. ***F***, Mitochondrial diameter comparison among subcompartments at P7 and P21. Box plot whiskers extend to the minimum/maximum within 1.5 interquartile range. Scale bars: ***A-E***, 5 µm.

Since we found that mitochondrial volumes in P21 calyces differ between axons and terminals (Fig. 6E), we further analyzed mitochondrial morphology by measuring the mitochondria diameter in axons (P7, P21), cup (P7), stalks and swellings (P21) from the same full resolution reconstruction of subcompartment data (Table 2). At P7, we found no difference between the mitochondrial diameters between axon and cup (0.4 ± 5 nm larger in the axon, *P* > 0.9999; Fig. 7F). Within subcompartments at P21 we found that stalks and swellings contained larger mitochondria diameters compared to the axon (44 ± 2 nm larger in stalks, *P* < 0.0001; 40 ± 2 nm larger in swellings, *P* < 0.0001), but found no difference between stalk and swellings (4 ± 2 nm larger in stalks, *P* = 0.1173). Across developmental stages, we observed an increase in the mitochondria diameter between P7 and P21 axons (21 ± 4 nm larger at P21, *P* < 0.0001) (Fig. 7F). Taken together, our findings reveal that stalks and swellings have similar mitochondrial volumes and diameters, and mitochondrial diameters are larger in P21 than in P7 calyces. Therefore, we conclude that differences in Ca^2+^ buffering between stalks and swellings are not caused by differences in mitochondrial volumes. Furthermore, the developmental increase in mitochondrial volumes is largely due to an increase in mitochondria size.

At P7, calyces had multiple filopodial extensions with various morphologies as shown in Figures 5A and 5D, some of which sparsely contained mitochondria (Fig. 6A, C). To further analyze the mitochondria distribution in these extensions we performed another full resolution partial reconstruction of a P7 calyx and overlaid its segmented mitochondria (Fig. 8, Movie 5). In this terminal, extensions from the cup-like calyx rarely contained mitochondria (Fig. 8B, D). In addition, we found multiple invaginations into the calyx from the principal cell, including large excrescences with complex membrane topologies (white arrows, Fig. 8A, C). These excrescences were largely composed of principal cell processes, though some processes could be traced back to the calyx terminal (Movie 5). Similar to filopodial extensions, these structures lacked mitochondria. This indicates that at P7 mitochondria are mostly restricted to the cup.

**Figure 8.**
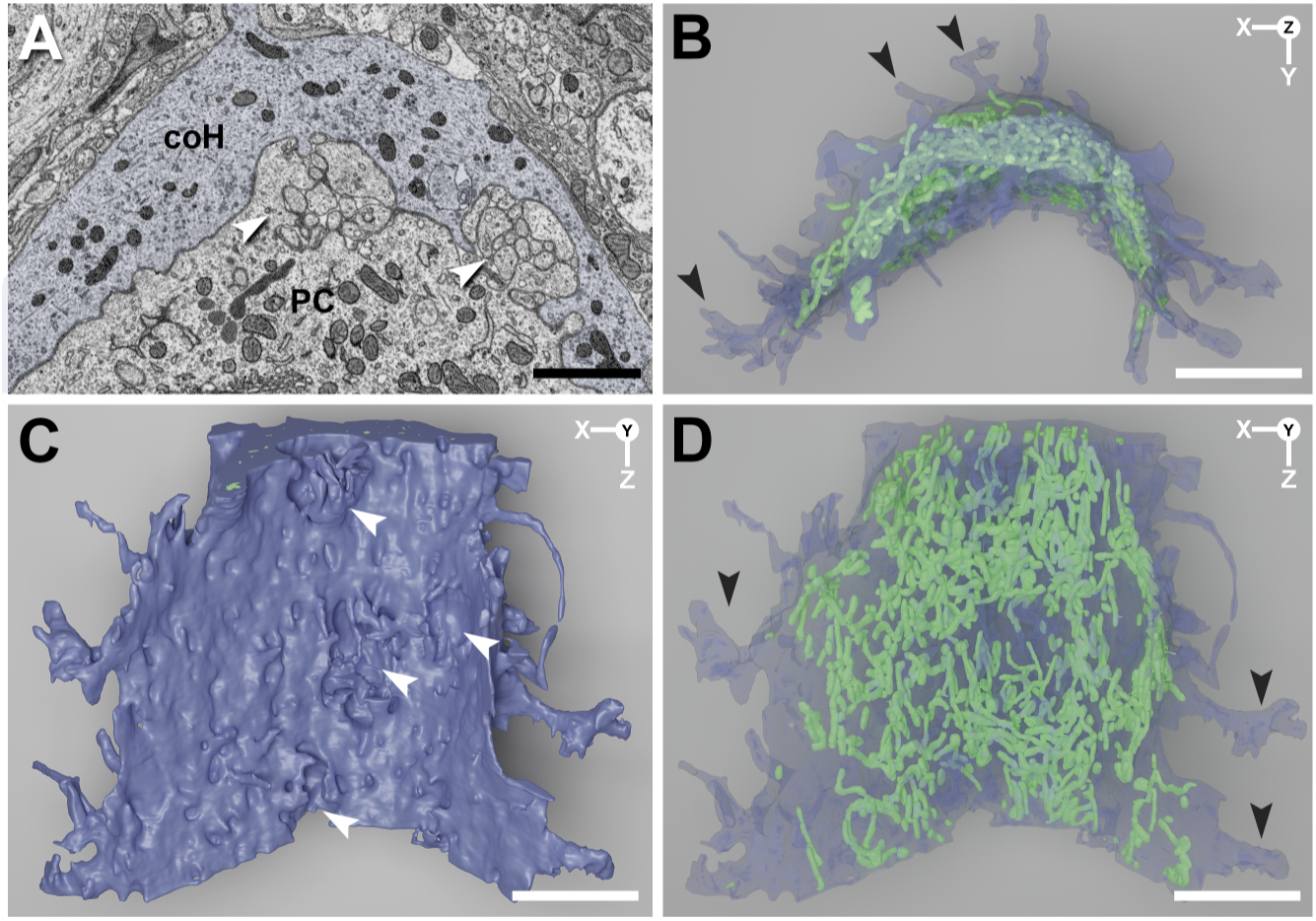
Mitochondria and membrane surface details of a segment of mito-APEX2-positive calyx of Held terminal reconstructed at full resolution from an SBF-SEM dataset at P7. ***A***, Part of a block face image of the terminal (blue) showing the organization of electron dense mitochondria. The principal cell-contacting face of the calyx terminal revealed a complex membrane topology. Protrusions of the principal cell invade the calyx, visualized as indentations in the calyx membrane. White arrows highlight excrescences of the principal cell (also shown in ***C***). ***B***, Top down view of the reconstructed terminal (blue) with mitochondria (green). Mitochondria are generally excluded from filopodial extensions of the calyx (black arrows in ***B*** and ***D***). ***C***, ***D***, Face-on view of terminal surface (***C***) and mitochondria (green) within the terminal (***D***), which form a complex network. Scale bars: ***A***-***D*** 5µm.

### Mito-APEX2 in conjunction with ATUM ssSEM allows for fine scale 3D analysis of presynaptic ultrastructure

In addition to using SBF-SEM in combination with mito-APEX2 to analyze mitochondrial volumes, one of our goals was to use mito-APEX2 as a genetic tool to analyze presynaptic ultrastructure in a defined population of genetically labeled neurons. However, we were unable to reliably detect the AZ with SBF-SEM. Therefore, we used ATUM-ssSEM to obtain SEM images with higher signal to noise ratio. We first established protocols to reliably detect mito-APEX2-labeled mitochondria with ATUM ssSEM and then used mito-APEX2 to carry out partial reconstructions of multiple AZs in the calyx.

Using our protocols, we could clearly identify labeled mitochondria in calyx terminals with detailed synaptic structure after post-staining sections to improve signal-to-noise ratio (SNR) for better membrane visualization in both P7 (Fig. 9A, B) and P21 (Fig. 9C, D, see Methods). Imaging with ssSEM was performed with several sessions at progressively higher resolutions. Beginning with a resolution of 2 µm/pixel, we collected images covering each section on a wafer, which required roughly a few hours. Following this, we imaged a broad region of interest in the middle of the MNTB containing dozens of principal cells at 30 nm/pixel resolution and captured ∼500 images to discriminate mito-APEX2-labeled terminals and to identify potential sites containing the fewest section wrinkles. Finally, we captured high resolution images at 5 nm/pixel resolution and a pixel dwell time of ∼13 µs, which was sufficient for analysis of synaptic specializations (Fig. 9B, D top). We morphologically differentiated AZs (Fig. 9B, D, top) from puncta adherentia (PA) (Fig. 9B, D, bottom) and separately quantified them for each calyx. Similar to SBF-SEM, we confirmed that calyx terminals containing mito-APEX2 positive mitochondria could be differentiated in both P7 and P21 ATUM-ssSEM samples (Fig. 9E, F).

**Figure 9.**
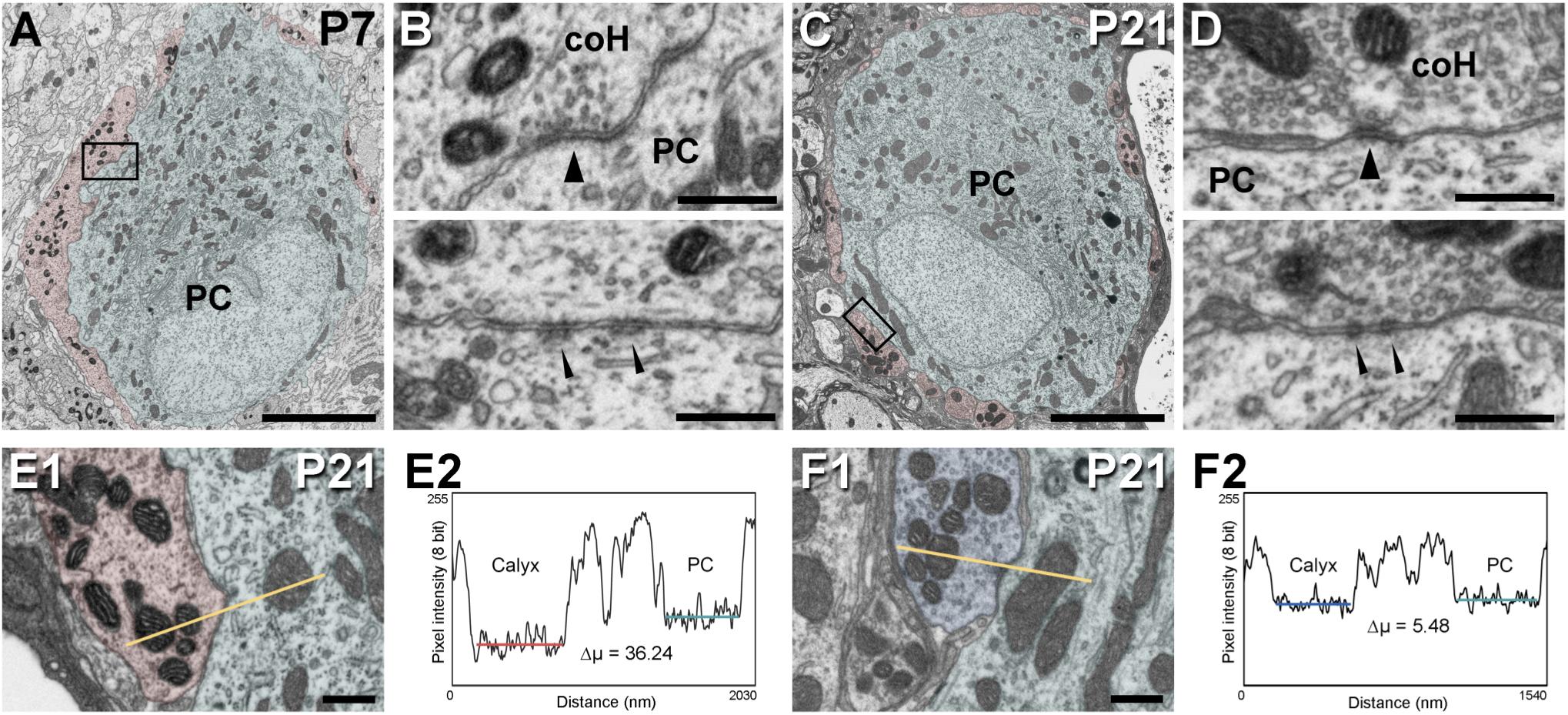
Mito-APEX2 labeling visualized by ssSEM at P7 (***A***, ***B***) and P21 (***C, D, E1***) allows for identification of transduced calyx of Held terminals. ***A***, ***C***, Ultrathin sections show calyx of Held input (red) to an MNTB principal cell (aqua) at P7 and P21, respectively. ***B, D***, Zoomed view of the terminals (top) depicted in ***A*** or ***C***, respectively, showing electron dense mitochondria and an active zone (top, thick arrowheads). Puncta adherentia (PA; bottom, thin arrowheads) are also located at the calyx-principal cell interface. High signal-to-noise ratio by ssSEM imaging allows for discrimination of individual SVs as well as pre- and post-synaptic densities. ***E1***, Transduced terminals contain darker mitochondria compared to mitochondria of the opposing principal cell. ***F1***, Non-transduced terminal mitochondria compared to the principal cell mitochondria. ***E2***, ***F2***, Intensity line profiles of mitochondrial intensity in transduced and non-transduced terminals, Δµ = difference in mean pixel intensity between calyx and PC mitochondria respectively. coH, calyx of Held; PC, principal cell; Scale bars: ***A***, ***C***, 5 µm; ***B***, ***D***, ***E***, ***F***, 500 nm.

Next, using 100 high-resolution serial sections captured from both P7 and P21 samples expressing mito-APEX2, we created partial reconstructions and carried out volumetric analysis for two calyces at each age (Fig. 10). The estimated surface areas of the partially reconstructed calyces facing the soma ranged from 117 to 269 µm^2^. (Fig. 10A, B, Table 2, and Movie 6 and 7). At P7, the number of PAs was approximately two-fold higher than AZs (37% AZ vs. 63% PA; Fig. 10A, Table 2), similar to previously reported values in P9 rat (Satzler et al., 2002). At P21 we observed a similar proportion of 40% AZ to 60% PA (Fig. 10B, Table 2).

**Figure 10.**
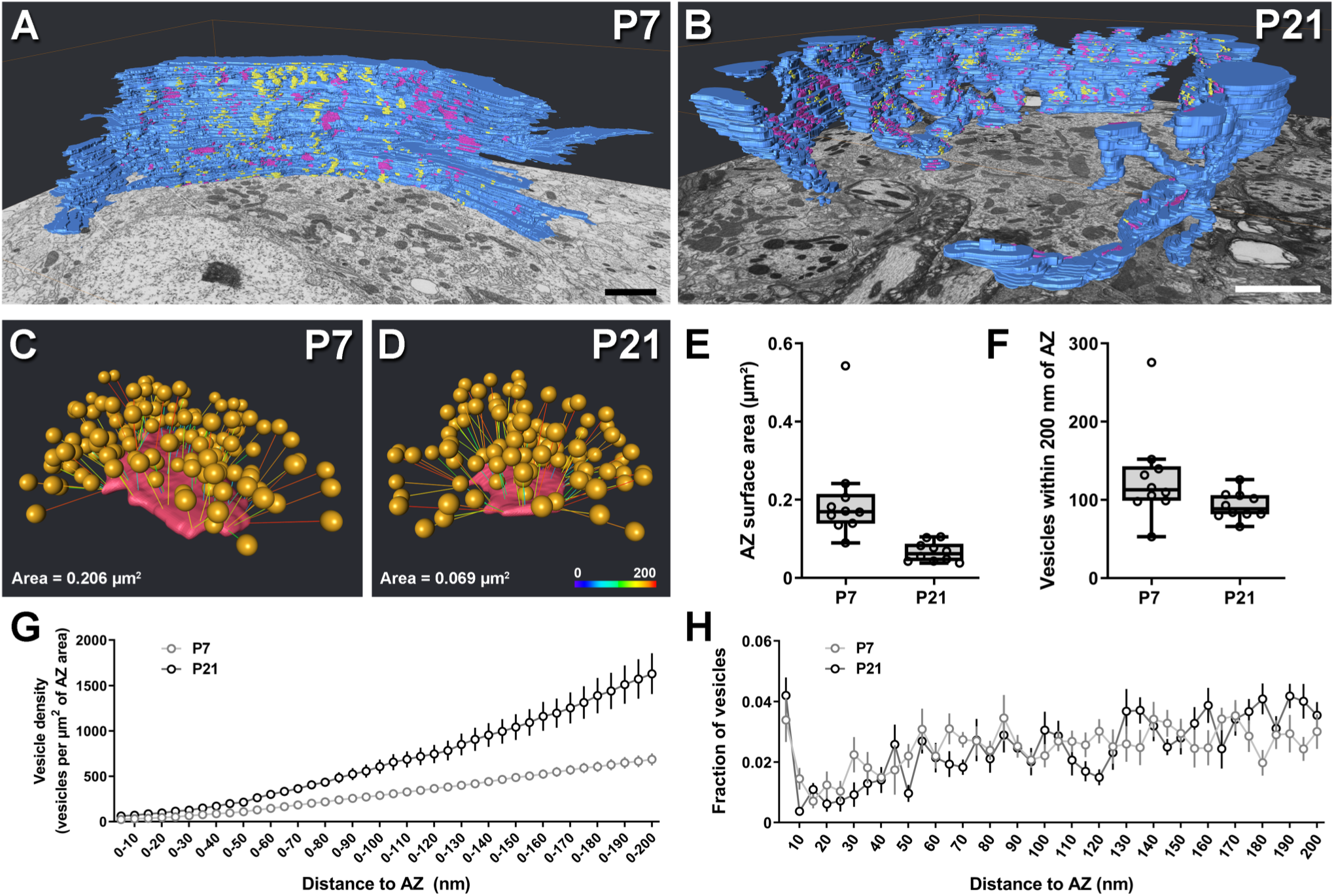
Identification of transduced terminals by ssSEM for 3D synaptic analysis. ***A***, ***B***, Partial 3D reconstructions from 100 serial sections of the calyx of Held terminals at P7 (***A***) and P21 (***B***) respectively, showing active zones (pink) and puncta adherentia (yellow). ***C***, ***D***, Ball models of an example active zone (pink) at the calyx of Held-MNTB synapse. SVs within 200 nm of the presynaptic membrane are modeled in 3D as 46 nm spheres (orange). Distances to the AZ from 0 to 200 nm are represented as colored lines. ***E***, AZ surface area at P7 and P21. ***F***, Total number of vesicles within 200 nm of the AZ. ***G***, Vesicle density as a function of AZ area, binned from 0 to 200 nm from the AZ. ***H***, Fraction of SVs within 5 nm bins up to 200 nm from the AZ. Box plot whiskers extend to the minimum/maximum within 1.5 interquartile range, open circles indicate individual observations. coH, calyx of Held; PC, principal cell; Scale bars: ***A***, ***B***, 5 µm; ***C D***, colorimetric scale bar 0-200 nm.

Since both AZ size and SV distributions change during early development (Taschenberger et al., 2002), we analyzed AZ area and SV distributions of 10 individual AZs (5/calyx) from 3D reconstructed models at P7 and P21. We found larger AZ at P7 compared to P21 (P7: 0.203 ± 0.040 µm^2^, n = 10 vs. P21: 0.066 ± 0.008 µm^2^, n = 10; Fig 10C, D, E, and Table 2). Comparing SV distribution within 200 nm of the AZ we found that AZ at P7 contained more SVs than P21 (P7: 128.2 ± 18.55, n = 10; vs. P21: 92.9 ± 5.363, n = 10; Fig. 10F, G, Table 2). However, when normalizing the SV number to AZ area to correct for the difference in AZ size, we found that the P21 AZ contained more SVs within 200 nm per AZ unit area (Fig. 10G). We observed no difference in overall SV distribution between P7 and P21 in a 200 nm distance from AZs in 3D space (Fig. 10H). At both ages, docked SVs (<5 nm to AZ) per AZ were similar (Table 2) and comparable to previous reports (Satzler et al., 2002; Taschenberger et al., 2002; Dondzillo et al., 2010), however we did not observe a developmental decrease in contrast to a previous study (Taschenberger et al., 2002).

## Discussion

To interrogate how mitochondria volumes are regulated during the development of the calyx of Held presynaptic terminal we developed a new tool, HdAd mito-APEX2, and optimized protocols and visualization workflows for use with SBF-SEM. We found that during development mitochondrial volumes increase in both the calyx and its respective axon, but that the mature calyx contains a larger relative mitochondrial volume than the axon. In addition, there was no difference in the relative mitochondrial volumes in the stalk and swellings of the mature calyx. Therefore, we propose that mitochondria levels are developmentally increased at the calyx of Held and its respective axon to support the energetic demands of high-frequency firing required to encode auditory information but do not significantly contribute to the calyx’s morphological-functional diversity. Finally, we believe as a technical advancement, that our HdAd vector in combination with optimized protocols for ATUM-SEM and SBF-SEM will be of broad use in combination with molecular perturbations studies using LM and EM inside and outside of the neuroscience community.

### A tubular network of mitochondria throughout development

Whole-terminal volumetric reconstructions of mitochondria at the two developmental stages revealed a complex tubular network in the P7 and P21 calyx. These findings of many long finger-like mitochondria are in line with previous reports from partial calyx reconstructions from P8 and P22 rats (Wimmer et al., 2006) and adult cat (Rowland et al., 2000) using ultra-thin section TEM. Long tubular mitochondria have been observed in multiple neuron types and their respective axons and across species using different imaging modalities, EM or fluorescence microscopy in both live cell and fixed tissues (Brodin et al., 1999; Misgeld et al., 2007; Saxton and Hollenbeck, 2012; Rossi and Pekkurnaz, 2019). However, the extensive tubular network might be specific to large synapses with high energy demands since many conventional presynaptic terminals in the CNS have a mitochondrion at or near the presynaptic terminal (Devine and Kittler, 2018). Since mitochondria can be functionally coupled (Amchenkova et al., 1988; Saxton and Hollenbeck, 2012), the tight packing and interconnected mitochondria observed in this and previous studies (Wimmer et al., 2006) suggest that mitochondria in the calyx and many other large presynaptic terminals are functionally coupled.

### Mitochondria become larger during calyx development

Based on our finding that mitochondria are more densely packed and mitochondria volumes are higher in the mature calyx compared to immature calyx, we propose that a developmental increase in mitochondrial volumes is critical for high-frequency transmission. Qualitative analysis of P7 calyces revealed large patches of the calyx without mitochondria while this was not found at the P21 calyx. Quantitative analysis of mitochondria diameters revealed that mitochondrial diameters were larger at the P21 calyces than those at P7. Though we found larger mitochondria at the mature stage, an increase in the total number of mitochondria may also contribute to the increase in relative volume. Obtaining absolute counts of mitochondria is difficult due to their ability to fuse and divide. However, qualitatively it appears that P21 calyces also contain more mitochondria. Although spontaneous activity in the auditory brainstem increases during early development (Tritsch and Bergles, 2010), the increase in mitochondrial volumes was larger at the calyx (∼3-fold), compared to the secondary inputs (∼1.4-fold) on the MNTB principal neuron. However, at P7 minor terminals had higher relative mitochondrial volumes (∼2.5-fold) compared to the calyx.

What are the signals that promote this transformation? During calyx development, synaptic activity increases ∼10-fold which lead to high rates of SV release and replenishment (Baydyuk et al., 2016). Thus, a probable signal is synaptic activity which is closely linked to mitochondria function (Nguyen et al., 1997; Rossi and Pekkurnaz, 2019). Total mitochondrial volume correlates with levels of exocytosis (Ivannikov et al., 2013), while individual mitochondrial volumes are larger at high-activity synapses (Cserep et al., 2018). In addition, mitochondria are more likely to pause during high activity levels and reside in the same position over a time course of days (Obashi and Okabe, 2013). Therefore, it possible that the increase in calyx activity increases the residency time of mitochondria in the terminal.

Comparison of axonal mitochondria volumes revealed that mitochondria were also more densely packed in P21 compared to P7. This indicates that increases in axonal excitability during development (Leao et al., 2005) are supported by increased mitochondria number to meet the energy demands of AP firing (Harris et al., 2012). Unlike the P7 axon, the mature calyx axon is myelinated except for the nodes of Ranvier and the heminode. Na^+^ channels are highly clustered at the heminode and excluded from the terminal (Leao et al., 2005). Although the last heminode in the mature calyx axon is well-defined using confocal imaging (Berret et al., 2016; Xu et al., 2017), we had difficulty defining the heminode from the myelination morphology only. Despite this, we were unable to detect any local increases in mitochondrial densities along the last 5-50 µm of the axons in the reconstructed volume. Therefore, our data indicate that mitochondria levels in the axon are globally increased without higher densities at the heminode. However, since mitochondrial movement is on the order of seconds (Devine and Kittler, 2018) it is possible that mitochondria pause movement at these heminodes during AP firing to create temporary increases in density which may not be detectable due to the timescale of aldehyde fixation.

We also found that mitochondrial volumes in the mature calyx were greater than its respective axon, while this was not found in the P7 calyx. This finding suggests SV release and replenishment in the mature calyx has higher energetic demands than AP firing, while the energetic demands at P7 maybe equivalent between the calyx and its respective axon. The mitochondrial adherens complex (MAC) is located within the mature calyx but has not been reported in the axon (Rowland et al., 2000). Therefore, it is possible that the MAC contributes to the increase in mitochondrial volume at the mature calyx due to its tethering of mitochondria. However, due to the limitations of SBF-SEM we were unable to identify the MAC and determine if the complex appearance of mitochondria in the calyx is developmentally regulated. Future studies will need to be carried out to investigate this further.

### Mitochondria and their contribution to the calyx morphological-functional continuum

Although the calyx is considered a reliable relay synapse (Englitz et al., 2009; Lorteije et al., 2009; Stasiak et al., 2018) a morphological-functional continuum of calyces that differ in the number of stalks and swellings in the hearing animal has been observed (Grande and Wang, 2011; Wang et al., 2015).
These differences in calyx morphologies corresponded to different short-term plasticity phenotypes, with simple calyces being strongly depressing and highly fenestrated calyces having initial facilitate with less depression in response to train stimuli (Grande and Wang, 2011). Differences in Ca^2+^ buffering and *Pr* between stalks and swellings has been proposed to expand the coding capacity of the calyx/MNTB synapse (Fekete et al., 2019). Mitochondria contribute to Ca^2+^ buffering (Billups and Forsythe, 2002; Guo et al., 2005; Verstreken et al., 2005) which can impact SV release (Devine and Kittler, 2018). In particular at the prehearing calyx, mitochondria slow the clearance of Ca^2+^ which can impact sustained synaptic transmission (Billups and Forsythe, 2002). However, at the prehearing calyx, mitochondria contribution to synaptic transmission is only prominent when the Na^+^/Ca^2+^ exchanger is saturated (Kim et al., 2005). Although swellings contain larger and faster Ca^2+^ transients and more Cav2.1 channel clusters (Fekete et al., 2019), we found no difference in the relative mitochondria volumes between stalks and swellings. Thus, it is possible that the contribution of mitochondria to the morphological-functional continuum may be minimal. However, we did not carry out functional analysis to perturb mitochondria and determine the impact on synapses in stalks and swelling. In addition, it is possible that only under long periods of high firing rates would mitochondria contribute to difference in Ca^2+^ clearance in these subcompartments. Future studies will need to be carried out to further investigate if mitochondrial play a role in the morphological-functional continuum.

### Identification of multiple inputs at the principal cells of the MNTB

Consistent with a previous report (Hoffpauir et al., 2006), we found multiple inputs innervating the MNTB principal neuron at P7. Strikingly, at P21 we also found labeled minor terminals in addition to a single calyx terminal. Although injection of HdAd with the human synapsin promoter labels other cell types in the CN in addition to the globular bushy cells (GBCs, (Montesinos et al., 2011), none of these other cell types are reported to make synapses on MNTB principal cells (Cant and Benson, 2003). Therefore, these labeled terminals are likely of GBC origin and distinct from local excitatory and inhibitory inputs (Hamann et al., 2003; Albrecht et al., 2014). Due to limited volumes imaged in this study we were unable to traces these axons to their origin. Since GBC axons give rise to multiple collaterals which form other small terminals (coincident) onto MNTB cells (Morest, 1968; Rodriguez-Contreras et al., 2006; Rodriguez-Contreras et al., 2008) it is possible that they are the source of the minor terminals that we observed. Future studies will need to be carried out to determine their role in regulating AP firing in the MNTB principal neuron.

### HdAd Mito-APEX2 as presynaptic marker for morphological studies

HdAds are a dsDNA viruses and have faster onset of transgene expression than single-stranded rAAVs which require second-strand synthesis (Hastie and Samulski, 2015; Montesinos et al., 2016). We took advantage of this fast onset of expression and carried out the EM analysis as early as P7, six days after HdAd injection, to analyze immature calyx ultrastructure. Although we did not test earlier time points, HdAd vectors have led to successful transduction of calyces as early as 72 hours (Montesinos et al., 2011). Since HdAd expression allows for long-term expression of transgenes for years (Kim et al., 2001) and our data demonstrate lack of toxicity of synapsin promoter driven expression of EGFP and mito-APEX2, this suggest that use of HdAd EGFP-mito-APEX2 will enable long term morphological studies on the scales of months to possibly years. More importantly, our HdAd vectors allows for the labeling of specific populations of neurons. When using mitochondria for volumetric reconstruction, the cellular compartments of interest should have a high mitochondria density for reliable distinction from unlabeled cells or compartments.

### ATUM ssSEM vs. SBF-SEM for presynaptic ultrastructure analysis

Use of HdAd mito-APEX2 with ATUM ssSEM allowed us to analyze presynaptic ultrastructure in addition to mitochondrial volumes. Both SBF-SEM and ATUM ssSEM have their strengths and weaknesses, therefore the different visualization platforms should be selected based on the target ultrastructure. Although SBF-SEM permitted large scale volumetric analysis, we were unable to segment fine AZ structures, such as the MAC or the puncta adherentia due to the low SNR required for consistent sectioning. Additionally, the low SNR prevented analysis of SV distribution. The low SNR was a result of the minimum acceptable beam dosage required to avoid sample damage and achieve consistent cutting of the block. Future improvements will need to be developed to enable better SNR in combination with thinner cutting thickness for fine structure analysis. Since ATUM ssSEM allows longer dwell times and therefore better SNR than SBF-SEM it was ideal for measuring SV distributions and AZ areas in a relatively large terminal. However, the tradeoff is the inherent issue with section alignment and possible compression artifacts, thus ATUM ssSEM is ideal for small scale volumetric reconstructions. Using ATUM ssSEM we found that the SV density per AZ area increases during development. While a previous study found a decrease in docked SV number from P5 to P14 (Taschenberger et al., 2002), we observed no change between P7 and P21. However, we only analyzed 10 AZs at each stage and differences in fixation protocols may lead to this discrepancy. Taken together, we believe that our viral vector and methods will be broadly applicable to quantitative analyses of subsynaptic organization of SVs, mitochondria, and their morphological continuum using TEM, ssSEM and SBF-SEM.

## Acknowledgements

We would like to thank the members of the Young lab for their comments on the manuscript, Keisuke Ohta for initial discussions, Melissa A. Ryan and Michael Morehead for help with EM volume data analysis. This work was supported by the National Institute of Deafness and Communication Disorders (R01 DC014093), the University of Iowa, Max Planck Society (SMY) and a German Research Foundation postdoctoral fellowship (DFG 420075000) (CK).

## Movie legends

**Movie 1.** Whole reconstructions of calyx of Held terminals contacting six MNTB principal cells at P7 in a 78 by 78 by 52 µm volume using SBF-SEM. Related to Figure 5A.

**Movie 2.** Whole reconstructions of calyx of Held terminals contacting six MNTB principal cells at P21 in a 78 by 78 by 79 µm volume using SBF-SEM. Related to Figure 5B.

**Movie 3.** Mitochondria reconstructions of calyx of Held terminals contacting six MNTB principal cells at P7 in the same volume as Movie 3. Related to Figure 6A.

**Movie 4.** Mitochondria reconstructions of calyx of Held terminals contacting six MNTB principal cells at P21 in the same volume as Movie 4. Related to Figure 6B.

**Movie 5.** Semi-automated partial reconstruction of a calyx of Held terminal at P7 in a 23 by 22 by 15 µm volume using SBF-SEM. Mitochondria (orange) were also semi-automatically segmented from the volume. Related to Figure 8.

**Movie 6.** Partial reconstruction of a labeled calyx of Held terminal at P7 in a 26 by 20 by 5 µm volume using ssSEM. Active zones (pink) and puncta adherentia (yellow) are shown on the calyx face in contact with the MNTB principal cell. Related to Figure 10A.

**Movie 7.** Partial reconstruction of a labeled calyx of Held terminal at P21 in a 27 by 21 by 5 µm volume using ssSEM. Active zones (pink) and puncta adherentia (yellow) are shown on the calyx face in contact with the MNTB principal cell. Related to Figure 10B.

